# GPR43 in eosinophils prevents the emergence of pathogenic Siglec-F^hi^ neutrophils in allergic airway inflammation

**DOI:** 10.1101/2024.08.07.607109

**Authors:** Jihyun Yu, Seongryong Kim, Hyun-Sup Song, Hye-Young Kim, You-Me Kim

**Author notes:** Corresponding author: You-Me Kim.

## Abstract

Eosinophils are major effector cells in type 2 immune responses, contributing to host defense and allergic diseases. They also play critical roles in maintaining tissue homeostasis by regulating various immune cell types. However, evidence of the crosstalk between eosinophils and neutrophils is limited. Here, we show that eosinophils directly associate with neutrophils in the lungs of asthma-induced mice. Eosinophil-specific deficiency of the short-chain fatty acid receptor GPR43 results in hyperactivation of eosinophils and increases the expression of neutrophil chemoattractants and PECAM-1, thus enhancing the interaction between eosinophils and neutrophils. This binding event exposes the neutrophils to eosinophil-derived IL-4 and GM-CSF, which induces the conversion of conventional neutrophils into more pathogenic Siglec-F^hi^ neutrophils that strongly enhance Th17 cell differentiation and aggravate asthma symptoms. These results reveal GPR43 as a critical regulator of eosinophils and highlight that eosinophils have a hitherto little-known ability to directly modulate neutrophil differentiation and function.

**One Sentence Summary:** Eosinophils directly recruit neutrophils and induce their differentiation into a pathogenic Siglec-F^hi^ subset in allergic airway inflammation.

## INTRODUCTION

Eosinophils are multifunctional granulocytes that are primarily recruited during type 2 immune responses. Once activated, they degranulate and release pre-synthesized cationic proteins, cytokines, chemokines, and lipid mediators to protect the host from infectious agents. However, this potent response can also cause tissue damage in allergic diseases (*1–3*). Moreover, eosinophils are constitutively present in several peripheral organs, where they maintain tissue homeostasis (*4, 5*). For example, eosinophils in adipose tissue are the major source of IL-4, which sustains the alternatively activated macrophages that control glucose metabolism (*6*). Intestinal eosinophils not only dominate the myeloid cells in the gastrointestinal tract but also prevent inflammation by suppressing Th1 and Th17 cells (*7, 8*). In addition, eosinophils in the lung protect it from allergen-induced airway inflammation by suppressing dendritic cells (DCs) (*9*). Notably, these local eosinophils markedly differ from newly infiltrating inflammatory eosinophils in terms of morphology and functions (*9*). Thus, eosinophils have diverse functions and make complex interactions with other immune cell types.

The development, migration, and activation of eosinophils are regulated by common β chain cytokines, such as IL-3, IL-5, and GM-CSF (*1, 10*). In addition, recent studies suggest that tissue environmental factors and nutrient-derived signals can control eosinophil functions. For example, aryl hydrocarbon receptor signaling in intestinal eosinophils induces transcriptional programming that permits their adaptation to the local environment, extends their life span, and confers their unique characteristics (*11, 12*). Similarly, the programming of intestinal eosinophils is also driven directly by retinoic acid: this signaling is specifically required for maintaining a villus-resident subpopulation of intestinal eosinophils (*13*). Moreover, the binding of neuromedin-U, a neuropeptide produced in the gut, to its receptor on eosinophils promotes their intestinal accumulation under normal conditions and induces goblet cell differentiation during helminth infection (*14*).

An interesting nutrient-derived factor is short-chain fatty acids (SCFAs), which are mainly produced by the gut microbiota via dietary fiber fermentation and circulated throughout the body (*15, 16*). SCFAs are recognized by GPR43, which is expressed by many non-immune and immune cells, including eosinophils (*17, 18*). Significantly, numerous studies show that SCFAs exert anti-inflammatory effects, and the lack of GPR43 signaling aggravates inflammatory diseases such as food allergy, colitis, rheumatoid arthritis, and asthma (*17, 19, 20*). Various GPR43-bearing cells have been found to mediate this anti-inflammatory activity, including DCs, neutrophils, type 3 innate lymphoid cells, and intestinal epithelial cells (*17, 19–22*). However, whether eosinophils also have GPR43-mediated anti-inflammatory activities is unknown.

Here, we showed with mice lacking GPR43 in eosinophils that GPR43 prevents eosinophils from becoming hyperactivated in allergic airway inflammation. Interestingly, this reduces direct eosinophil interactions with neutrophils in the lung, thereby suppressing the emergence of a pathogenic, Th17-promoting neutrophil subset. Thus, this study shows that GPR43 regulates eosinophil functions, and that eosinophils have a previously little-known ability to directly modulate neutrophil differentiation and activities in lung inflammation.

## RESULTS

### GPR43 deficiency in eosinophils exacerbates asthma in association with increased lung neutrophils and Th17 cells

GPR43 is encoded by free fatty acid receptor 2 *(FFAR2*). As previously observed (*14, 23–27*), *FFAR2* is constitutively expressed at high levels in eosinophils from various organs in humans and mice. By contrast, other *FFAR* genes (*FFAR1* and *FFAR3*) are poorly expressed (fig. S1, A and B). *FFAR2* is also strongly expressed in the lungs of asthmatic mice (fig. S1C). Moreover, publicly available single-cell RNA-sequencing (scRNA-seq) data from patients with eosinophilic esophagitis show that eosinophils in the inflamed esophagus and duodenum express *FFAR2* more strongly than other immune cell types, similar to the transcriptional profile of the eosinophil marker CCR3 (*17, 18*) (fig. S1, D and E). Given that GPR43 has well-known anti-inflammatory functions and eosinophils drive airway inflammation in asthma, we asked whether GPR43 regulates eosinophils in asthma.

To address this, GPR43 was specifically deleted in eosinophils by crossing *Gpr43*^fl/fl^ mice with *Epx-Cre* mice, thus generating *Epx*^Cre/+^*Gpr43*^fl/fl^ mice. Eosinophil-specific Cre-mediated recombination was confirmed with *Epx*^Cre/+^Rosa26-eYFP^fl/+^ reporter mice. As reported previously (*7*), YFP was exclusively expressed in eosinophils, of which 60-70% expressed YFP (fig. S2, A to C). Notably, despite the apparently incomplete recombination induced by *Epx-Cre* in the reporter mice, GPR43 mRNA was not detected in eosinophils from *Epx*^Cre/+^*Gpr43*^fl/fl^ mice (fig. S2D). Moreover, acetate, an endogenous GPR43 ligand, did not induce Ca^2+^ influx in eosinophils from *Epx*^Cre/+^*Gpr43*^fl/fl^ mice (fig. S2E). Thus, GPR43 signaling is efficiently abrogated in GPR43-deficient eosinophils.

*Epx*^Cre/+^*Gpr43*^fl/fl^ and control (*Epx*^Cre/+)^ mice were then subjected to allergic airway inflammation with house dust mite extracts (HDM) (Fig. 1A). Compared to *Epx*^Cre/+^ mice, *Epx*^Cre/+^*Gpr43*^fl/fl^ mice had significantly more immune cells, including eosinophils and neutrophils, in the bronchoalveolar space (Fig. 1B). Histological analysis revealed more inflammation, mucus accumulation, and fibrosis in the lungs of *Epx*^Cre/+^*Gpr43*^fl/fl^ mice (Fig. 1C). Accordingly, *Epx*^Cre/+^*Gpr43*^fl/fl^ mice showed worse airway hyperresponsiveness compared to control mice (Fig. 1D). Thus, GPR43 deletion in eosinophils aggravated airway inflammation and asthma symptoms. The bronchoalveolar lavage fluid (BALF) from asthmatic *Epx*^Cre/+^*Gpr43*^fl/fl^ mice also contained higher type 2 cytokine levels (IL-4, IL-5, and IL-13) (Fig. 2A) and their lungs bore more GATA3- and type 2 cytokine-expressing CD4 T cells (Fig. 2, B to D). Notably, the BALF from *Epx*^Cre/+^*Gpr43*^fl/fl^ mice also contained significantly more IL-17A, and the lungs bore higher frequencies of RORγt- and IL-17A-expressing CD4 T cells in the lung (Fig. 2, B to D). These results show that GPR43 deficiency in eosinophils enhances both Th2 and Th17 immune responses in asthma. Similarly, GPR43 deletion in eosinophils induced more severe ovalbumin (OVA)-induced airway inflammation that was associated with enhanced Th17 responses in the lung (fig. S3, A to F).

**Fig. 1.**
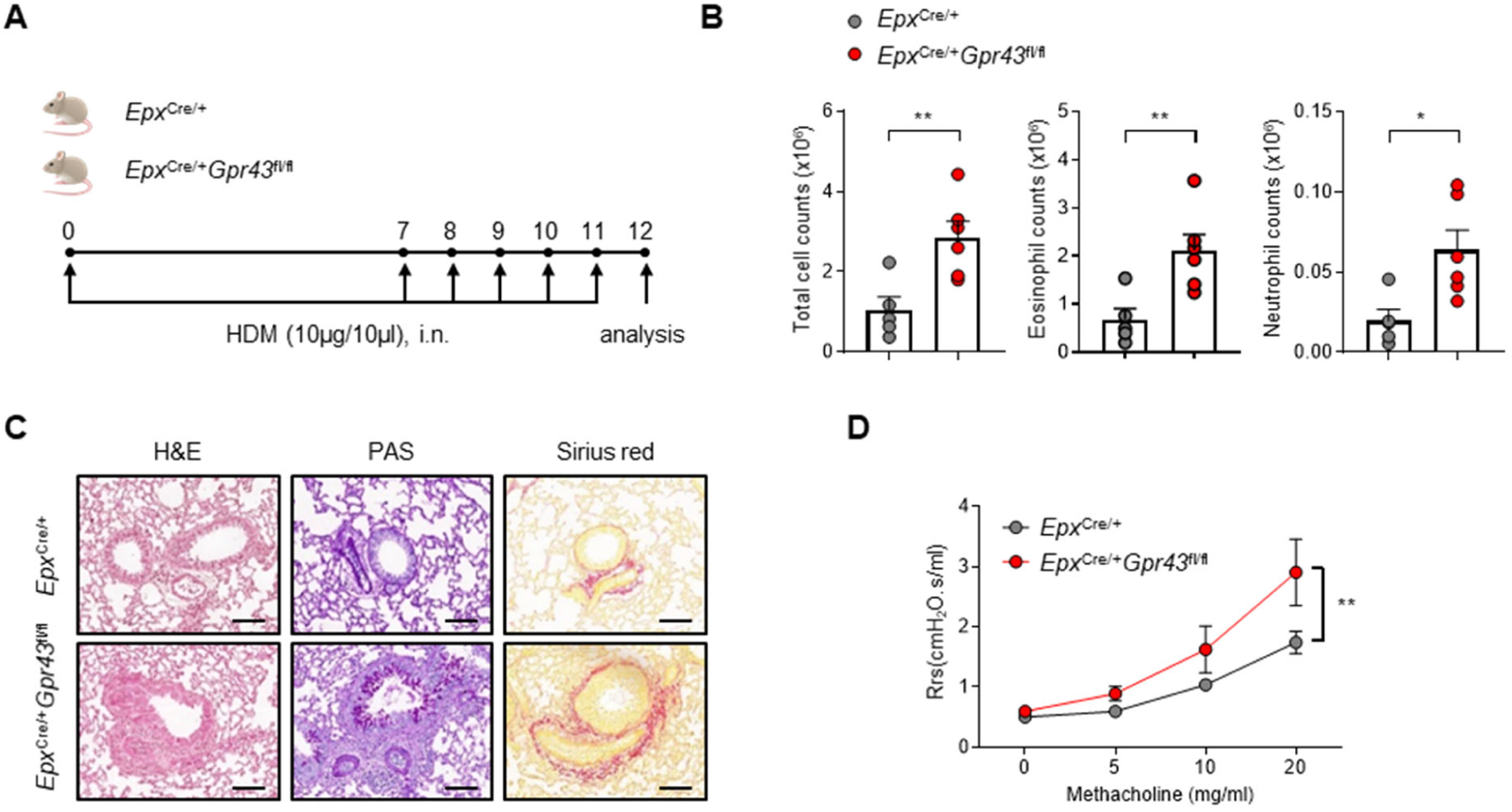
GPR43 deficiency in eosinophils exacerbates HDM-induced asthma. (**A**) Experimental scheme of HDM-induced asthma. (**B**) Total immune cell, eosinophil, and neutrophil count in BALF. *Epx*^Cre/+^ (n = 5), *Epx*^Cre/+^*Gpr43*^fl/fl^ (n = 6). Data are representative of more than three independent experiments. (**C**) Histological analysis of immune cell infiltration (H&E), mucus production (PAS), and fibrosis (Sirius red) in the asthmatic lungs. Scale bar: 100 µm. Data are representative of two independent experiments. (**D**) Airway resistance test. *Epx*^Cre/+^ (n = 9), *Epx*^Cre/+^*Gpr43*^fl/fl^ (n = 9). Data were pooled from two independent experiments. Data are presented as mean ± s.e.m. Unpaired two-tailed Student’s t-test (B) and two-way ANOVA (D) were performed for statistical analysis (* p < 0.05, ** p < 0.01).

**Fig. 2.**
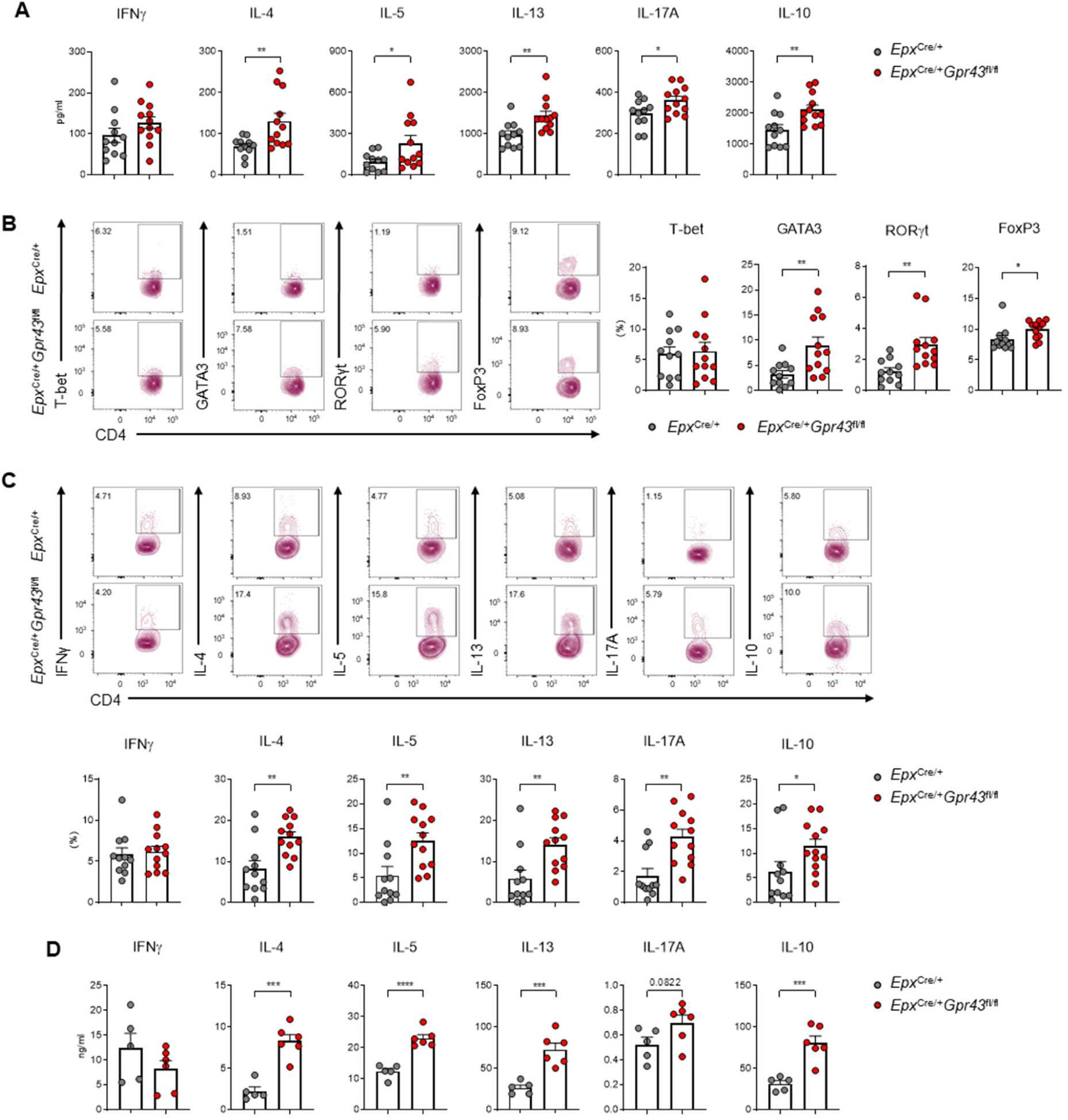
GPR43 deficiency in eosinophils upregulates Th2 and Th17 responses in HDM-induced asthma. (**A**) Cytokine levels in BALF. (**B**) The percentage of CD4 T cells expressing each transcription factor in asthmatic lungs. (**C**) The percentage of CD4 T cells producing each cytokine in asthmatic lungs. *Epx*^Cre/+^ (n = 11), *Epx*^Cre/+^*Gpr43*^fl/fl^ (n = 12). Data in (A-C) were pooled from two independent experiments. (**D**) Cytokine levels in the culture supernatant of asthmatic lung cells stimulated with anti-CD3 and anti-CD28. *Epx*^Cre/+^ (n = 5), *Epx*^Cre/+^*Gpr43*^fl/fl^ (n = 6). Data in (D) are representative of three independent experiments. Data are presented as mean ± s.e.m. Unpaired two-tailed Student’s t-test was performed for statistical analysis (* p < 0.05, ** p < 0.01, *** p < 0.001, **** p < 0.0001).

### Pathogenic Siglec-F^hi^ neutrophils accumulate in the asthmatic lungs of *Epx*^Cre/+^*Gpr43*^fl/fl^ mice

To comprehensively examine the effect of eosinophil-specific GPR43 deletion on the lung immune cell landscape during airway inflammation, CD45^+^ cells from HDM-induced asthmatic lungs of *Epx*^Cre/+^ and *Epx*^Cre/+^*Gpr43*^fl/fl^ mice were subjected to scRNA-seq. Unsupervised cell clustering revealed nine groups bearing conventional immune cell type markers (Fig. 3A and fig. S4A). Notably, eosinophils were not detected. This was also observed in publicly available scRNA-seq datasets of healthy and asthmatic lungs (*27*, *28*) and probably reflects high RNase levels in eosinophils (*28–30*). The most notable difference between *Epx*^Cre/+^ and *Epx*^Cre/+^*Gpr43*^fl/fl^ mice was the neutrophils: they were highly enriched in *Epx*^Cre/+^*Gpr43*^fl/fl^ mice (Fig. 3B). Fine clustering of T cells also confirmed that GPR43 deletion in eosinophils increased *Gata3-*, *Il4-*, *Il5-*, and *Il13-*expressing Th2 cells and *Rorc-* and *Il17-*expressing Th17 cells (fig. S4, B to D).

**Fig. 3.**
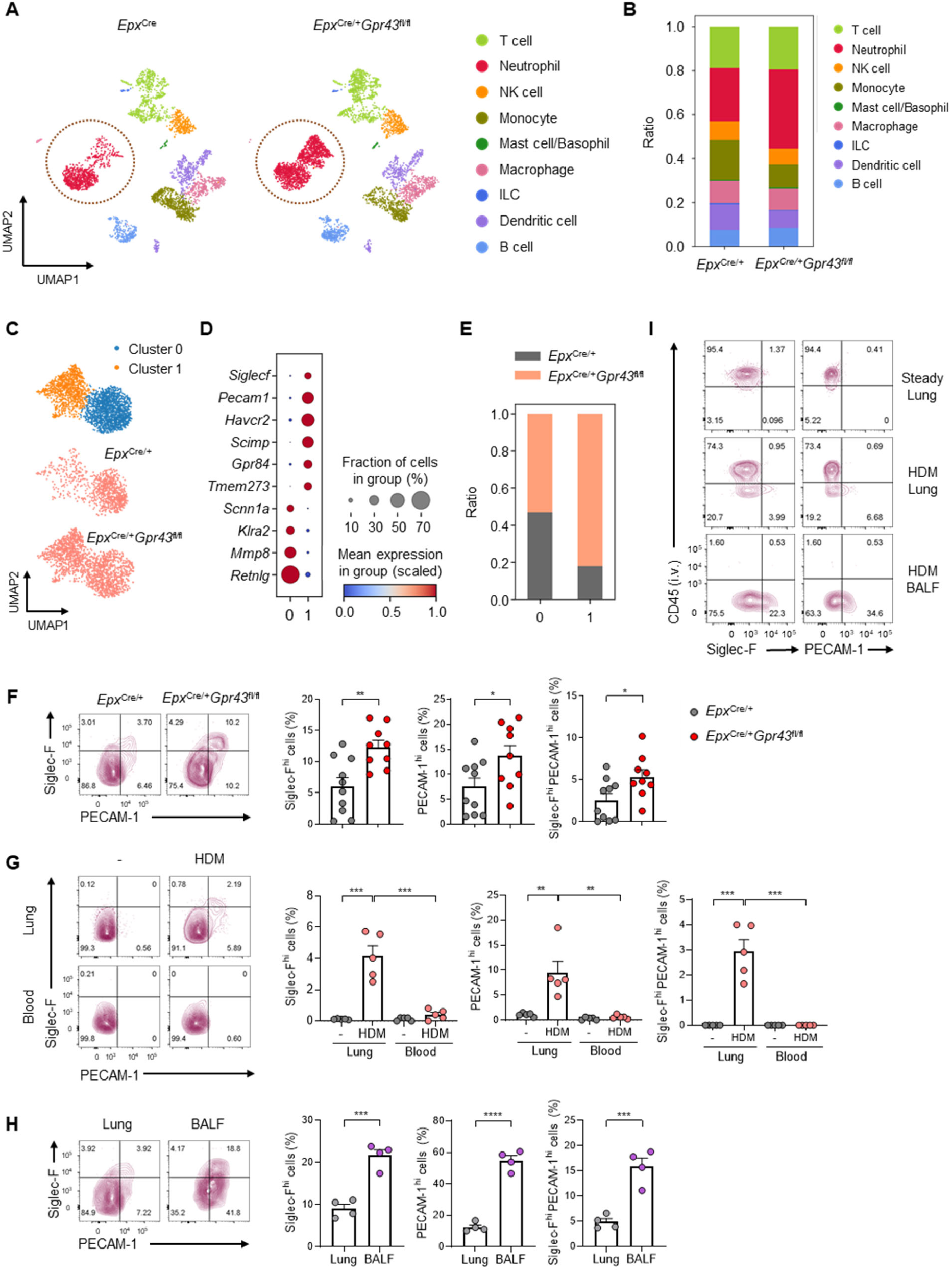
Absence of GPR43 in eosinophils results in the emergence of pathogenic Siglec-F^hi^ neutrophils in asthmatic lungs. (**A**) UMAP plots of immune cells in asthmatic lungs of *Epx*^Cre/+^ and *Epx*^Cre/+^*Gpr43*^fl/fl^ mice. (**B**) Ratio of the immune cell types in each group. (**C**) UMAP plots of neutrophils showing two distinct clusters. (**D**) Dot plot for marker genes of each neutrophil cluster. (**E**) Ratio of each group in the individual neutrophil clusters. (**F**) The percentage of Siglec-F^hi^ and Siglec-F^hi^PECAM-1^hi^ cells among neutrophils in asthmatic lungs. *Epx*^Cre/+^ (n = 10), *Epx*^Cre/+^*Gpr43*^fl/fl^ (n = 9). Data were pooled from two independent experiments. (**G**) The percentage of Siglec-F^hi^ and Siglec-F^hi^PECAM-1^hi^ cells among neutrophils in the lung and blood of steady-state and asthmatic mice. *Epx*^Cre/+^ mice (n = 5). (**H**) The percentage of Siglec-F^hi^ and Siglec-F^hi^PECAM-1^hi^ cells among neutrophils in the lung and BALF of asthmatic mice. *Epx*^Cre/+^ mice (n = 4). (**I**) Staining of neutrophils in the steady-state lung, asthmatic lung, and asthmatic BALF with intravenously administered anti-CD45 antibodies. Data in (G-I) are representative of three independent experiments. Data are presented as mean ± s.e.m. Unpaired two-tailed Student’s t-test was performed for statistical analysis (* p < 0.05, ** p < 0.01, *** p < 0.001).

Subclustering of all neutrophils in our scRNA-seq datasets resulted in two distinct clusters (Fig. 3, A and C). Both expressed *Cxcr2* and *S100a8* but other genes typically associated with neutrophils (e.g. *Mmp8* and *Retnlg*) demonstrated dramatic downregulation in Cluster 1. Moreover, several other genes were almost exclusively expressed in Cluster 1, namely, *Siglecf*, *Pecam1*, *Havcr2*, *Scimp*, *Gpr84*, and *Tmem273* (Fig. 3D and fig. S5A). Siglec-F was particularly interesting because, while it often serves as an eosinophil marker, recent studies showed that Siglec-F-expressing neutrophil subsets associate with various diseases such as lung adenocarcinoma, myocardial infarction, and diesel exhaust particle (DEP)-induced lung inflammation (*31–33*). In the latter, Siglec-F^hi^ neutrophils actively produce neutrophil extracellular traps (NETs) and promote the accumulation of type 2 cytokine-producing cells in the lung (*33*). Moreover, the sputum of asthma patients bears neutrophils that express Siglec-8 (a human paralog of murine Siglec-F), and Siglec-8^+^ neutrophil frequencies in patient peripheral blood correlate positively with asthma severity (*33*). Thus, the Siglec-F-expressing neutrophils in the asthmatic lungs from *Epx*^Cre/+^*Gpr43*^fl/fl^ mice may be pathogenic. Notably, the transcriptomic data from lung adenocarcinoma, myocardial infarction, and DEP-induced lung inflammation studies (*31–33*) showed that many of the genes that were upregulated in their Siglec-F^hi^ neutrophils (relative to Siglec-F^low^ neutrophils) were also highly enriched in our Cluster 1 neutrophils (fig. S5B). Therefore, we designated our Cluster 1 and Cluster 0 neutrophils as Siglec-F^hi^ and Siglec-F^low^ neutrophils, respectively. Cluster 0 (Siglec-F^low^) neutrophils were composed of the neutrophils from both *Epx*^Cre/+^ and *Epx*^Cre/+^*Gpr43*^fl/fl^ mice at approximately 1:1 ratio, but most Cluster 1 (Siglec-F^hi^) neutrophils were from *Epx*^Cre/+^*Gpr43*^fl/fl^ mice (Fig. 3E). This enrichment of Siglec-F^hi^ neutrophils in asthmatic lungs of *Epx*^Cre/+^*Gpr43*^fl/fl^ mice was confirmed by flow cytometry (Fig. 3F), and was also observed in the OVA-induced asthma model (fig. S3G). Thus, while some Siglec-F^hi^ neutrophils were present in asthmatic lungs of control mice, they were greatly increased when GPR43 was deleted in eosinophils.

Siglec-F^hi^ neutrophils were found only in asthmatic lungs: the healthy lungs or peripheral blood of healthy and asthmatic mice only bore Siglec-F^low^ neutrophils (Fig. 3G). Additionally, the frequencies of Siglec-F^hi^ neutrophils were significantly higher in the BALF than in the lungs (Fig. 3H). Thus, Siglec-F^hi^ neutrophils likely differentiate from Siglec-F^low^ neutrophils in the lung during airway inflammation and infiltrate the airway more efficiently. This was supported by *in vivo* labeling of circulating neutrophils with intravenously injected CD45 antibodies just before sacrifice: none of Siglec-F^hi^ neutrophils in the asthmatic lungs and BLAF were labeled with anti-CD45 antibodies. Thus, Siglec-F^hi^ neutrophils are exclusively localized in the lung tissue and exudate, but not in its microvasculature (*34*) (Fig. 3I).

### GPR43-deficient eosinophils directly induce the differentiation of Siglec-F^hi^ neutrophils

We next asked how the lack of GPR43 in eosinophils caused the emergence of Siglec-F^hi^ neutrophils during airway inflammation. First, we compared control (EOS^Ctrl^) and GPR43-deficient eosinophils (EOS^△GPR43^) from the asthmatic lungs of *Epx*^Cre/+^ and *Epx*^Cre/+^*Gpr43*^fl/fl^ mice, respectively. EOS^△GPR43^ were highly vacuolized and exhibited disorganized nuclear shapes (Fig. 4, A to C). Eosinophils from patients with severe asthma also have these features (*35*), which suggests that EOS^△GPR43^ were hyperactivated. The transcriptomic analysis by bulk-RNA sequencing revealed that activated eosinophils expressed many genes related to neutrophil chemotaxis, several of which were expressed at higher levels in EOS^△GPR43^ than in EOS^Ctrl^ (fig. S6, A and B). Thus, we assessed the ability of the eosinophils to directly attract neutrophils. Indeed, when EOS^Ctrl^ or EOS^△GPR43^ were cultured with bone marrow-derived neutrophils in a transwell system (Fig. 4D), neutrophil migration was enhanced by both eosinophils but especially by EOS^△GPR43^ (Fig. 4, E and F). Moreover, unlike neutrophils cultured alone, neutrophils co-cultured with EOS^Ctrl^ or EOS^△GPR43^ formed aggregates, especially with EOS^△GPR43^ (Fig. 4G). Immunofluorescence imaging of the aggregates revealed direct contact between neutrophils and eosinophils, especially with EOS^△GPR43^ (Fig. 4, H and I). Immunofluorescence imaging of the asthmatic lung of *Epx*^Cre/+^ mice also showed physical interactions between the neutrophils and eosinophils, especially around the airways (fig. S6C). Thus, eosinophils in the asthmatic lung actively attract and make contact with neutrophils, and this interaction is augmented by GPR43 deficiency in eosinophils. Notably, reanalysis of publicly available transcriptomic data showed that when peripheral blood eosinophils from asthma patients were activated *in vitro* with IL-33, many neutrophil chemotaxis-related genes were upregulated, implying that human eosinophils can also recruit neutrophils (fig. S6D).

**Fig. 4.**
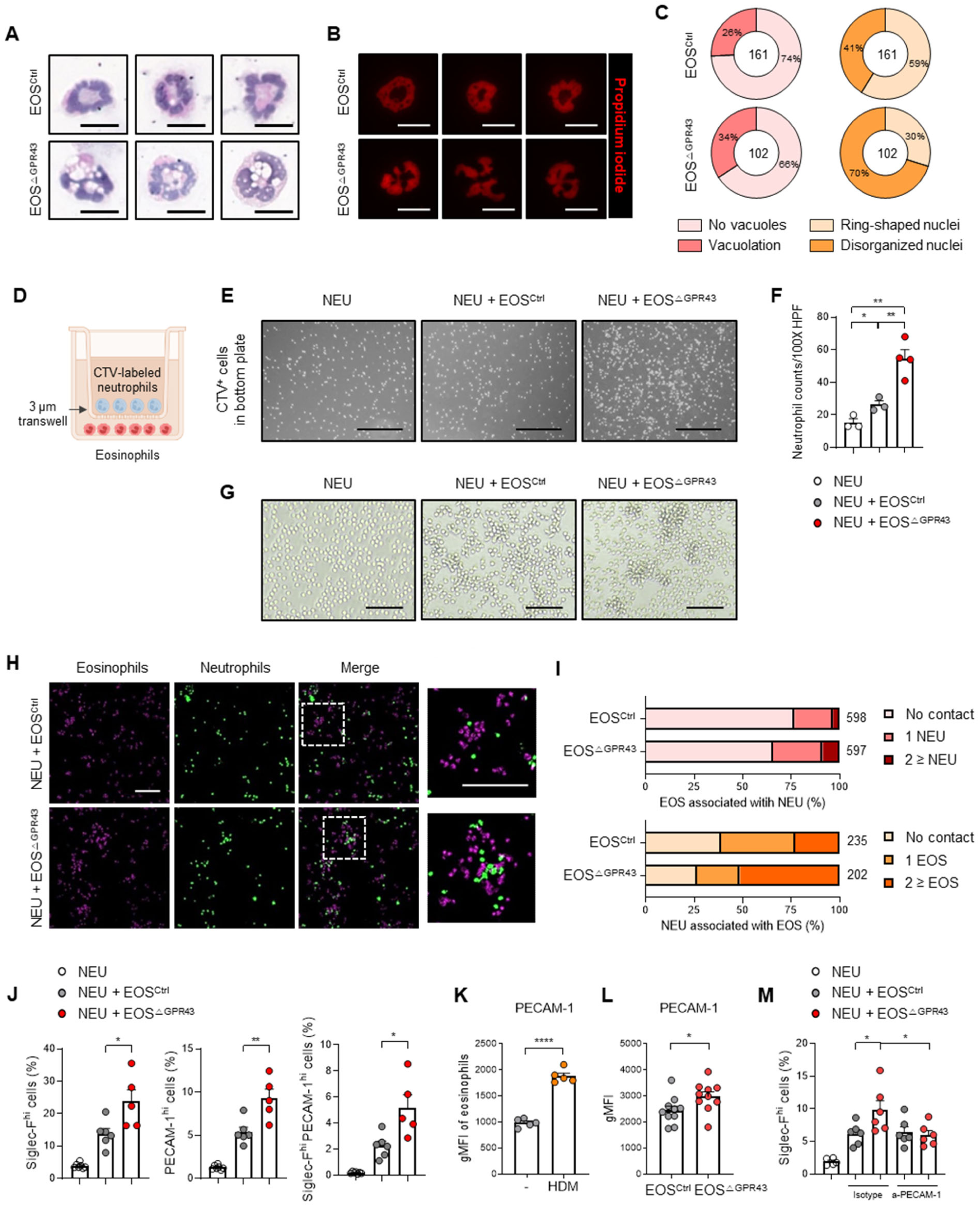
GPR43-deficient eosinophils attract neutrophils more efficiently and promote the differentiation of Siglec-F^hi^ neutrophils. (**A**) Representative images of H&E-stained asthmatic lung eosinophils from *Epx*^Cre/+^ and *Epx*^Cre/+^*Gpr43*^fl/fl^ mice. Scale bar: 10 µm. (**B**) Representative images of propidium iodide-stained asthmatic lung eosinophils. Scale bar: 10 µm. (**C**) The percentage of asthmatic lung eosinophils showing vacuolation (left) and disorganized nuclei (right). The percentages were calculated from three independent H&E images for each group. The total numbers of eosinophils were indicated in the center of the donut plots. (**D**) Neutrophil migration assay scheme. (**E**) Representative images showing CTV-labeled neutrophils migrated toward eosinophils in the bottom plate. Scale bar: 200 µm. (**F**) Neutrophil counts within 100-fold magnified HPF images of the bottom plates. Asthmatic lung eosinophils were pooled from three mice per group for sorting, and assays were performed in technical replicates. NEU (n = 3), NEU + EOS^Ctrl^ (n = 3), NEU + EOS^△GPR43^ (n = 4). (**G**) Images of eosinophil-neutrophil co-culture. Scale bar: 100 µm. Data in (D-G) are representative of three independent experiments. (**H**) Immunofluorescence images of eosinophil-neutrophil co-culture. Scale bar: 100 µm. Data are representative of two independent experiments. (**I**) The percentage of eosinophils associated with neutrophils (top) and neutrophils associated with eosinophils (bottom). The percentages were calculated from four and three independent fluorescence images for the EOS^Ctrl^ and EOS^△GPR43^ groups, respectively. (**J**) The percentage of Siglec-F^hi^ and Siglec-F^hi^PECAM-1^hi^ cells among neutrophils after co-culture with asthmatic lung eosinophils. EOS^Ctrl^ (n = 6), EOS^△GPR43^ (n = 5). Data in (J) are representative of three independent experiments. (**K**) gMFI of PECAM-1 in eosinophils from steady-state (n = 5) and asthmatic lungs (n = 5). Data are representative of three independent experiments. (**L**) gMFI of PECAM-1 in asthmatic lung eosinophils. *Epx*^Cre/+^ (n = 10), *Epx*^Cre/+^*Gpr43*^fl/fl^ (n = 10). Data were pooled from two independent experiments. (**M**) The percentage of Siglec-F^hi^ and Siglec-F^hi^PECAM-1^hi^ cells among neutrophils after co-culture with eosinophils in the presence or absence of anti-PECAM-1. EOS^Ctrl^ (n = 6), EOS^△GPR43^ (n = 5). Data are representative of three independent experiments. Data in (J-M) are biological replicates. Data are presented as mean ± s.e.m. Unpaired two-tailed Student’s t-test (F and J-M) were performed for statistical analysis (* p < 0.05, ** p < 0.01, **** p < 0.0001).

The physical association between eosinophils and neutrophils in asthmatic lungs led us to ask whether eosinophils directly caused Siglec-F^low^ neutrophils to differentiate into Siglec-F^hi^ neutrophils. Thus, bone marrow-derived neutrophils, which are Siglec-F^low^, were co-cultured with eosinophils from asthmatic or healthy lungs of control mice for 2 days. In the absence of eosinophils, few Siglec-F^hi^ neutrophils arose. However, this differentiation was increased by steady-state lung eosinophils, and hugely elevated by eosinophils from asthmatic lungs (fig. S6E). Moreover, the latter enhancement was further augmented by GPR43 deficiency in eosinophils (Fig. 4J). Thus, the eosinophils in the asthmatic lung likely directly drove the emergence of Siglec-F^hi^ neutrophils, and GPR43 deletion in eosinophils worsened it.

Notably, Siglec-F^hi^ neutrophils from asthmatic lungs also demonstrated upregulation of the *Pecam1* gene (Fig. 3D and fig. S5A), and a large proportion of Siglec-F^hi^ neutrophils in the asthmatic lung and BALF co-expressed PECAM-1 protein (Fig. 3, F to H). Similar to Siglec-F^hi^ neutrophils, PECAM-1^hi^ neutrophils were only detected in the lung and BALF, but not in blood, after asthma induction (Fig. 3, G and H). Eosinophil-specific deletion of GPR43 also elevated PECAM-1^hi^ and Siglec-F^hi^PECAM-1^hi^ neutrophil frequencies (Fig. 3F). Similarly, many Siglec-F^hi^ neutrophils generated from bone marrow-derived neutrophils by co-culture with EOS^Ctrl^ or EOS^△GPR43^ also co-expressed PECAM-1, especially with EOS^△GPR43^ (Fig. 4J). Interestingly, we found that asthma induction upregulated PECAM-1 expression not only on neutrophils but also on eosinophils (Fig. 4K). This eosinophil PECAM-1 expression was higher on EOS^△GPR43^ than on EOS^Ctrl^ (Fig. 4L). PECAM-1 is an adhesion molecule that can mediate the binding of neutrophils to other cells in a homophilic manner (*36, 37*). Given the physical interactions between eosinophils and neutrophils *in vitro* (Fig. 4, G to I) and *in vivo* (fig. S6C), we speculated that PECAM-1 on lung neutrophils and eosinophils may bind together, thereby attaching the two cell types. To determine whether these homophilic interactions participate in eosinophil-induced Siglec-F^hi^ neutrophil differentiation, EOS^△GPR43^ or EOS^Ctrl^ were co-cultured with bone marrow-derived neutrophils in the presence of PECAM-1-blocking antibody. The antibody reduced the ability of EOS^△GPR43^ to induce Siglec-F^hi^ neutrophil differentiation but did not inhibit EOS^Ctrl^-induced Siglec-F^hi^ neutrophil differentiation (Fig. 4M). This suggests that (i) PECAM-1 proteins overexpressed on EOS^△GPR43^ critically contribute to the ability of these GPR43-deficient eosinophils to enhance Siglec-F^hi^ neutrophil differentiation, probably by promoting the tighter binding to neutrophils by EOS^△GPR43^ than EOS^Ctrl^, but (ii) there are still other mechanisms that mediate the interaction between eosinophils and neutrophils, thus making PECAM-1 dispensable for Siglec-F^hi^ neutrophil differentiation by EOS^Ctrl^.

Next, we sought to identify the eosinophil-derived mediators that promote Siglec-F^hi^ neutrophil differentiation. Thus, bone marrow-derived Siglec-F^low^PECAM-1^low^ neutrophils were stimulated with various cytokines or LPS. GM-CSF and, to a lesser extent, IL-33, TNF, and LPS, induced the neutrophils to express Siglec-F. By contrast, IL-4 and LPS, and to a lesser extent, IL-33, GM-CSF, and TGFβ, caused the neutrophils to express PECAM-1 (Fig. 5A). We then combined the cytokines that most strongly stimulated Siglec-F (GM-CSF) and PECAM-1 (IL-4) expression: this combination increased the frequencies of Siglec-F^hi^, PECAM-1^hi^, and Siglec-F^hi^PECAM-1^hi^ neutrophils in a highly synergistic manner (Fig. 5B). This efficient GM-CSF+IL-4-mediated induction of Siglec-F^hi^, PECAM-1^hi^, and Siglec-F^hi^PECAM-1^hi^ neutrophils was also observed when neutrophils from healthy lungs were treated with these cytokines (Fig. 5C). Notably, EOS^Ctrl^-induced differentiation of neutrophils into Siglec-F^hi^, PECAM-1^hi^, and Siglec-F^hi^PECAM-1^hi^ neutrophils was significantly inhibited by neutralizing antibodies against both IL-4 and GM-CSF (Fig. 5D). Thus, eosinophil-derived GM-CSF and IL-4 can drive Siglec-F^hi^-neutrophil differentiation and upregulate their PECAM-1 expression.

**Fig. 5.**
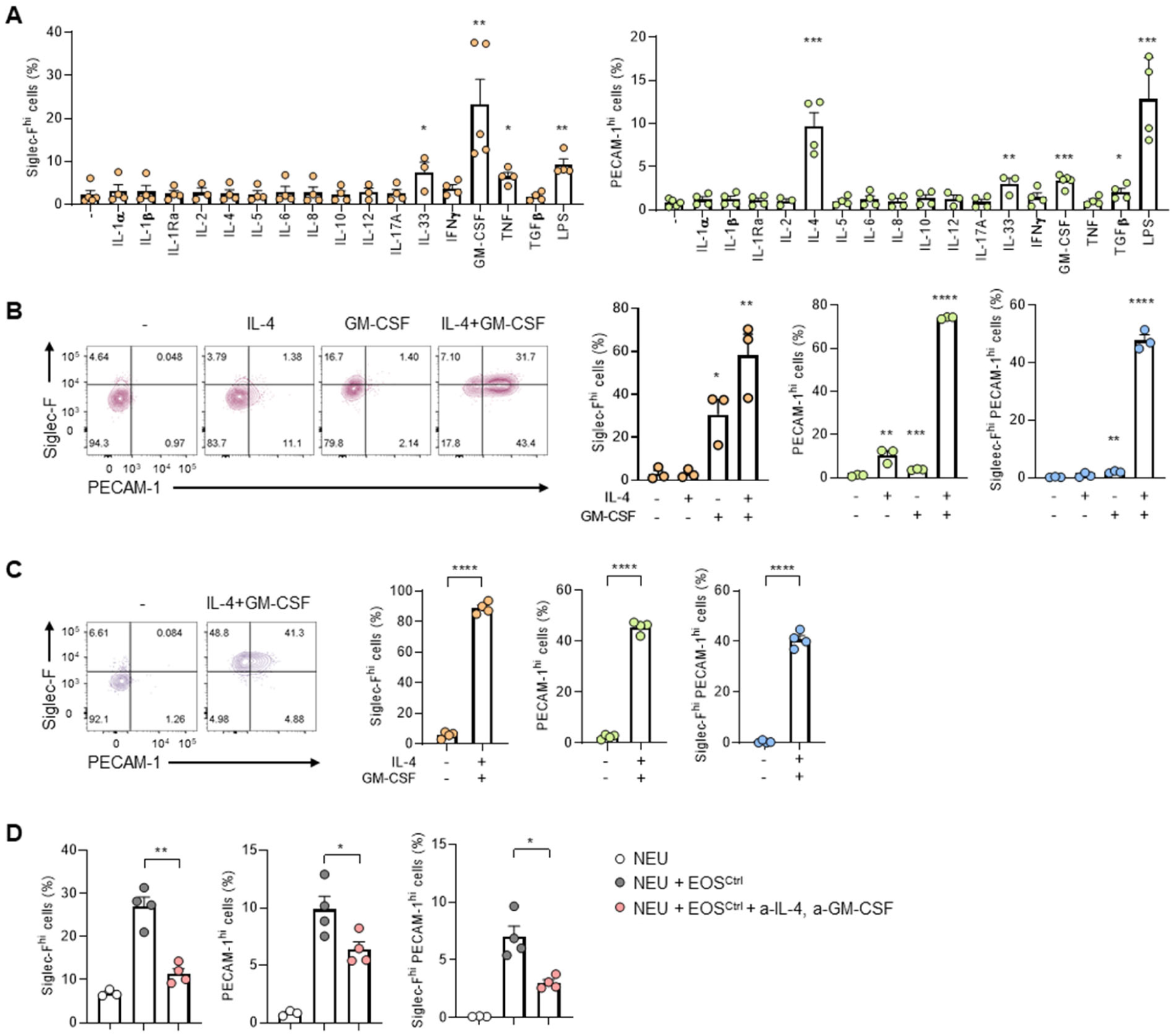
Eosinophil-induced differentiation of Siglec-F^hi^ neutrophils is dependent on IL-4 and GM-CSF. The percentage of Siglec-F^hi^, PECAM-1^hi^, or Siglec-F^hi^PECAM-1^hi^ neutrophils. (**A**) Bone marrow neutrophils were stimulated with various cytokines or LPS (n = 3-5). Data were pooled from five independent experiments. (**B**) Bone marrow neutrophils were stimulated with IL-4, GM-CSF, or both cytokines (n = 3). Data were pooled from three independent experiments. (**C**) Steady-state lung neutrophils were stimulated with the combination of IL-4 and GM-CSF (n = 4). Data are representative of two independent experiments. (**D**) Bone marrow neutrophils were co-cultured with asthmatic lung eosinophils in the presence or absence of anti-IL-4 and anti-GM-CSF antibodies (n = 4). Data are representative of three independent experiments. Data are presented as mean ± s.e.m. Unpaired two-tailed Student’s t-test (A, B) and paired two-tailed Student’s t-test (C, D) were performed for statistical analysis (* p < 0.05, ** p < 0.01, *** p < 0.001, **** p < 0.0001).

### Siglec-F^hi^ neutrophils enhance Th17 differentiation

To better characterize the Siglec-F^hi^ neutrophils in the HDM-induced asthmatic lung, Siglec-F^low^ and Siglec-F^hi^ neutrophil transcriptomes were compared. Relative to Siglec-F^low^ neutrophils, Siglec-F^hi^ neutrophils demonstrated (i) strong upregulation of cytokine (e.g. *Csf1*, *Tnf*, *Il1a*), chemokine (*Ccl3*, *Ccl4*), degranulation (*Cd63*), and lipid-mediator synthesis (*Ptgs1*, *Ltc4s*) genes, and (ii) downregulation of classical neutrophil function genes (*Lyz2*, *Mmp8*, *S100a8*, *S100a9*, *Ly6g*, *Retnlg*) and a gene related to homing to secondary-lymphoid organs (*Sell*) (Fig. 6A). Gene set enrichment analysis also showed that ‘cytokine activity’, ‘TNF signaling pathway’, and ‘NF-kappa B signaling pathway’ genes were highly enriched in Siglec-F^hi^ neutrophils, as were genes that relate to rheumatoid arthritis and Coronavirus disease-COVID-19 (Fig. 6B and fig. S7). Moreover, Siglec-F^hi^ neutrophils demonstrated enrichment in pattern recognition receptor pathways such as ‘NOD-like receptor signaling pathway’ and ‘C-type lectin receptor signaling pathway’ along with ‘Phagosome’, ‘Lysosome’, ‘Ribosome’, ‘Oxidative phosphorylation’, and ‘Apoptosis’ genes (fig. S7). Thus, compared to conventional Siglec-F^low^ neutrophils, Siglec-F^hi^ neutrophils were more activated and likely performed more pro-inflammatory functions.

**Fig. 6.**
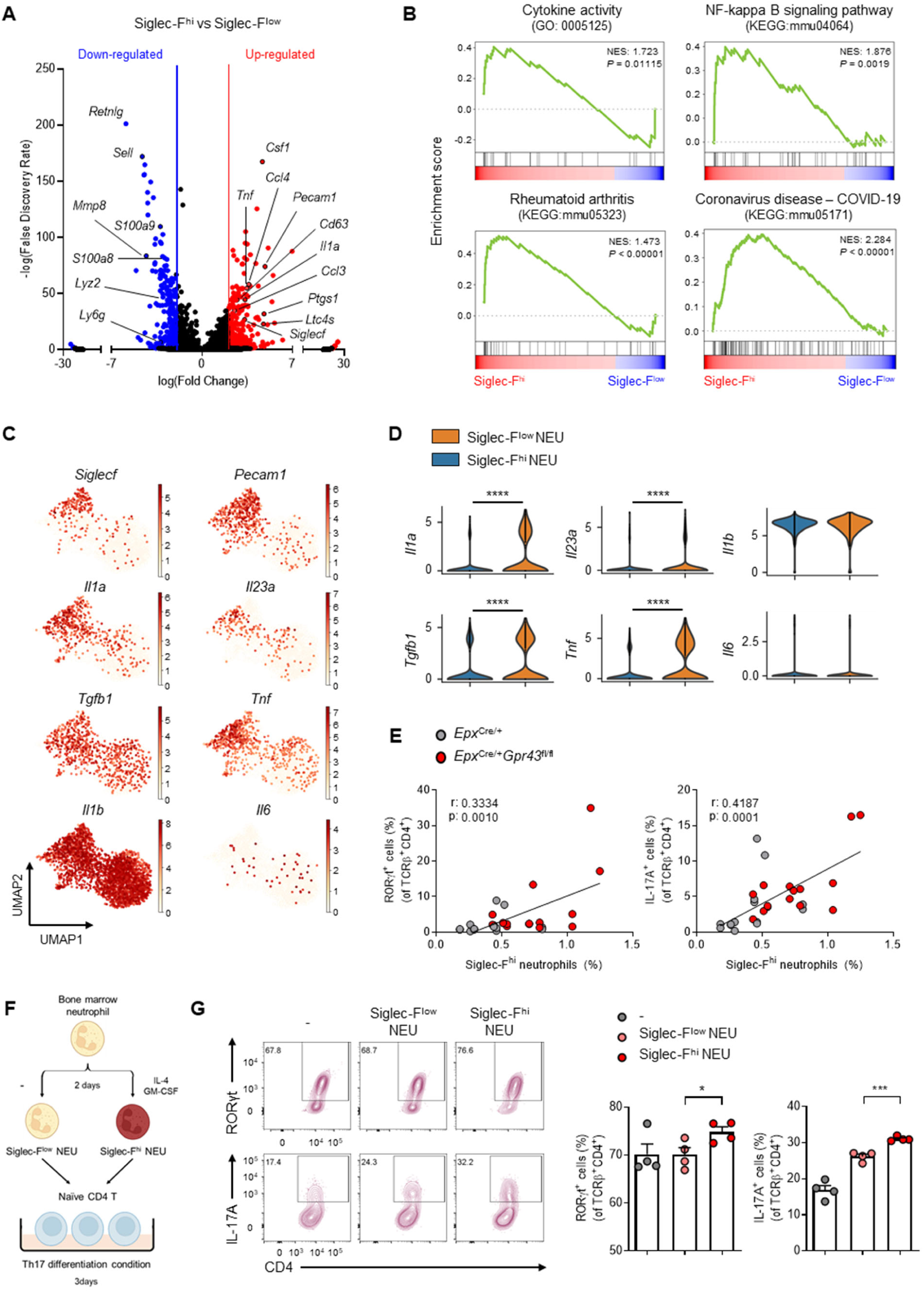
Siglec-F^hi^ neutrophils enhance the differentiation of Th17 cells. (**A**) Volcano plot displaying differential gene expression between Siglec-F^low^ and Siglec-F^hi^ neutrophils. Highlighted: FDR < 0.05, absolute logFC > 2. (**B**) Gene set enrichment analysis of DEGs between Siglec-F^low^ and Siglec-F^hi^ neutrophils. (**C**) UMAP plots and (**D**) violin plots for cytokine genes related to Th17 differentiation. (**E**) Correlation between the proportion of Siglec-F^hi^ neutrophils and RORγt^+^ or IL-17A^+^ CD4 T cells in asthmatic lungs. *Epx*^Cre/+^ (n = 14), *Epx*^Cre/+^*Gpr43*^fl/fl^ (n = 15). Data were pooled from three independent experiments. (**F**) Experimental scheme for co-culture of neutrophils and naïve CD4 T cells under Th17 differentiation condition. Siglec-F^low^ and Siglec-F^hi^ neutrophils were prepared by culturing bone marrow neutrophils in the absence or presence of IL-4 and GM-CSF, respectively. (**G**) The percentage of RORγt^+^ or IL-17A^+^ CD4 T cells after the co-culture. Siglec-F^low^ NEU (n = 4), Siglec-F^hi^ NEU (n = 4). Data are representative of four independent experiments. Data are presented as mean ± s.e.m. Simple linear regression (E) and unpaired two-tailed Student’s t-test (G) were performed for statistical analysis (* p < 0.05, *** p < 0.001, **** p < 0.0001).

Closer examination of the upregulated cytokines in Siglec-F^hi^ neutrophils revealed notable increases in Th17-promoting cytokines, namely, *Il1a*, *Il23a*, *Tgfb1*, and *Tnf*, although two other Th17-inducing cytokines were expressed at similar levels in Siglec-F^hi^ and Siglec-F^low^ neutrophils (*Il1b*) or were poorly expressed in both (*Il6*) (Fig. 6, C and D). Interestingly, of all immune cells in asthmatic lungs, neutrophils were largely the only producers of *Il1a*, *Il23a*, and *Tnf* (fig. S8A). Thus, Siglec-F^hi^ neutrophils may drive Th17 responses. Indeed, Siglec-F^hi^-neutrophil frequencies in asthmatic lungs correlated positively with RORγt^+^ and IL-17A^+^ CD4 T cell frequencies (Fig. 6E). To determine whether Siglec-F^hi^ neutrophils can directly drive Th17 differentiation, we generated Siglec-F^hi^ neutrophils from Siglec-F^low^ neutrophils with IL-4+GM-CSF (Fig. 6F). Like Siglec-F^hi^ neutrophils in asthmatic lungs, these *in vitro* generated Siglec-F^hi^ neutrophils expressed high levels of *Il1a, Il23a*, and *Tnf* compared to Siglec-F^low^ neutrophils (fig. S8B). They also expressed more IL-1α and TNF, as determined by ELISA (fig. S8C). Thus, Siglec-F^hi^ neutrophils generated *in vitro* with cytokines closely resembled Siglec-F^hi^ neutrophils in asthmatic lungs. Importantly, co-culture with naïve CD4 T cells showed that Siglec-F^hi^ neutrophils significantly enhanced Th17 cell differentiation compared to Siglec-F^low^ neutrophils, as indicated by higher RORγt^+^ and IL-17A^+^ cell frequencies (Fig. 6, F and G).

Eosinophils were shown to modulate CD4 T cell differentiation directly or indirectly through regulating DC functions (*7, 8, 38, 39*). We thus asked whether eosinophils can augment Th17 differentiation, like Siglec-F^hi^ neutrophils. However, co-culturing naïve CD4 T cells with EOS^Ctrl^ or EOS^△GPR43^ with or without DCs showed that while DCs increased Th17 differentiation, eosinophil could neither directly or indirectly enhance it (fig. S9, A and B). This is consistent with the comparable expression in EOS^Ctrl^ and EOS^△GPR43^ of antigen presentation- and Th17 differentiation-related genes (fig. S9, C and D) and their similar levels of IL-1-receptor antagonist (IL-1Ra), which can suppress Th17 cell differentiation (*8*) (fig. S9E). Moreover, DCs in asthmatic lungs from *Epx*^Cre/+^ and *Epx*^Cre/+^*Gpr43*^fl/fl^ mice demonstrated similar expression of antigen presentation-, Th17 activation-, and Th17 differentiation-related genes (fig. S9F). Collectively, these results suggest that lack of GPR43 in eosinophils induces their hyperreactivity in the asthmatic lung, which generates Siglec-F^hi^ neutrophils that then become the main drivers of pathogenic Th17 cells.

## DISCUSSION

GPR43 plays anti-inflammatory roles in various inflammatory diseases, and multiple cell types can mediate this activity. We show here that GPR43 also plays anti-inflammatory roles in eosinophils: eosinophils lacking GPR43 became hyperactive when allergic airway inflammation was induced. Our *in vitro* and *in vivo* observations suggest that this hyperactivity led to excessive eosinophil-induced differentiation of Siglec-F^hi^ neutrophils, which in turn increased Th17 cells in the lung, and more severe airway inflammation.

Eosinophils interact with and control diverse immune cell types. However, whether eosinophils can regulate neutrophils, particularly directly, has been poorly studied (*40, 41*), likely because eosinophils and neutrophils play canonical roles in type 2 and type 3 immunity, respectively (*42*). Our study showed that eosinophils can in fact directly regulate neutrophils in several ways. First, even normal lung eosinophils (with GPR43) from asthmatic mice could recruit neutrophils by producing neutrophil chemoattractants. Moreover, our re-analysis of publicly available bulk RNA-seq data showed that when human eosinophils are treated with the asthma-promoting cytokine IL-33, they express multiple neutrophil chemoattractants. This potential for direct neutrophil attraction by eosinophils has also been observed in other inflammatory settings (*43*). Second, we observed that eosinophils interacted physically with neutrophils, both *in vitro* and *in vivo*, and that this led to their differentiation into Siglec-F^hi^ neutrophils. Both mechanisms were elevated when eosinophils lacked GPR43, which suggests that GPR43 constitutively prevents eosinophils from recruiting neutrophils and converting them into more pathogenic Siglec-F^hi^ neutrophils.

The eosinophil-induced differentiation of Siglec-F^hi^ neutrophils was likely mediated by eosinophil-derived IL-4 and GM-CSF, since differentiation was largely blocked by antibodies to these cytokines. However, other factors may also contribute: previous studies show that ATP and TGFβ also promote Siglec-F^hi^ neutrophil differentiation (*33, 44*). We also found that IL-33 and TNF can generate Siglec-F^hi^ neutrophils, albeit less well than GM-CSF. Further identification of the signals required for *in vivo* generation of Siglec-F^hi^ neutrophils in allergic airway inflammation is needed.

We found that many Siglec-F^hi^ neutrophils in the asthmatic lungs and BALF co-expressed PECAM-1. Siglec-F^hi^PECAM-1^hi^ neutrophils were also generated *in vitro* from Siglec-F^low^PECAM-1^low^ neutrophils by asthmatic lung eosinophils, especially GPR43-deficient eosinophils. Notably, PECAM-1 co-expression by Siglec-F^hi^ neutrophils has not been observed elsewhere, including in lung adenocarcinoma (*31*), myocardial infarction (*32*), and DEP-induced lung inflammation (*33*). Since IL-4 was by far the most potent PECAM-1-upregulating cytokine of the 16 we tested, it is possible that PECAM-1 co-expression by Siglec-F^hi^ neutrophils may be unique to type 2 cytokine-rich inflammatory conditions. However, we found that LPS, GM-CSF, and IL-33 also induced PECAM-1 expression. Thus, PECAM-1 could potentially be upregulated in Siglec-F^hi^ neutrophils in other inflammatory conditions.

Since asthmatic lung eosinophils also express PECAM-1, and this is upregulated by GPR43 deletion, we speculated that homophilic PECAM-1 interactions induce binding of neutrophils to eosinophils and contribute to Siglec-F^hi^ neutrophil induction. However, PECAM-1-blocking antibody only suppressed the enhanced ability of EOS^△GPR43^ to induce Siglec-F^hi^ neutrophil differentiation: it did not suppress baseline EOS^Ctrl^-induced Siglec-F^hi^ neutrophil differentiation. Therefore, direct binding between eosinophils and neutrophils may not be a prerequisite for Siglec-F^hi^ neutrophil differentiation. Alternatively, other adhesion molecules play a more major role in the physical interaction between eosinophils and neutrophils, especially in the early phase of their interactions, while overexpressed PECAM-1 in EOS^△GPR43^ further enhances the interactions. Another possibility is that PECAM-1-mediated signaling events participate in the eosinophil-induced neutrophil differentiation: the cytoplasmic tail of PECAM-1 bears immunoreceptor tyrosine-inhibitory motifs, and PECAM-1 can modulate immune cell intracellular signaling (*37, 45*). Further studies are needed to determine the exact mechanism(s) by which PECAM-1 regulates eosinophil-induced Siglec-F^hi^-neutrophil differentiation.

Notably, asthmatic lungs also bore neutrophils that expressed PECAM-1 but not Siglec-F (Fig. 3F to G). Moreover, none of the PECAM-1^hi^ or Siglec-F^hi^ neutrophils were found in the circulation (Fig. 3I). In addition, genes related to pathogenic Th17 differentiation (*46, 47*) were upregulated in Siglec-F^hi^ neutrophils and they could enhance Th17 cell differentiation. These findings, together with the production of neutrophil chemoattractants by eosinophils in asthmatic lungs, suggest the following sequence of immunological events in asthmatic lungs: (1) PECAM-1^low^ neutrophils enter the lung from blood, partly due to eosinophil-derived chemokines, (2) the IL-4-rich local environment induces neutrophil expression of PECAM-1, (3) the eosinophil-derived chemokines further encourage the neutrophils to approach and physically connect with the eosinophils and the local eosinophil-derived IL-4 further increases PECAM-1 expression on the neutrophils, (4) eosinophil-derived IL-4+GM-CSF induces Siglec-F expression (i.e. differentiation) of the neutrophils, and (5) Siglec-F^hi^ neutrophils promote Th17 differentiation (fig. S10). This whole process can potentially occur in the asthmatic *Epx*^Cre/+^ lung but significantly worsens when GPR43 is deleted in eosinophils because the deletion increases eosinophil-mediated neutrophil chemoattraction and eosinophil PECAM-1 expression. Consequently, EOS^△GPR43^ attract more neutrophils and convert them to Siglec-F^hi^ neutrophils. Therefore, eosinophil-expressed GPR43 downregulates crosstalk between eosinophils and neutrophils in asthma-induced lung inflammation, thus minimizing eosinophil-mediated induction of Siglec-F^hi^ neutrophils.

Siglec-F^hi^ neutrophils have been found to play pathogenic roles in many diseases (*31–34*). Similarly, Siglec-F^hi^ neutrophils in our HDM-induced asthma model highly expressed proinflammatory genes, especially *Il1a* and *Il23a*, promoting pathogenic Th17 cells (*46, 47*). Since Th17 cells and IL-17A can, in turn, promote neutrophil-mediated immune responses (*48*), it is possible that there is a positive feedback loop between Siglec-F^hi^ neutrophils and Th17 cells that results in exacerbated asthma symptoms. Moreover, this feedback loop can be strengthened when GPR43 is deleted in eosinophils.

These findings are of clinical interest because asthma is broadly classified on the basis of immune cell profiles as eosinophilic and non-eosinophilic asthma (*49–51*). Eosinophilic asthma is driven by Th2 responses and is generally readily managed with corticosteroids. Non-eosinophilic asthma includes the neutrophilic and paucigranulocytic endotypes and associates with either Th1 or Th17 responses. It is often resistant to corticosteroid treatment. Non-eosinophilic asthma also includes the mixed-granulocytic endotype, which is characterized by Th2 and Th17 responses and joint eosinophil and neutrophil involvement: this endotype is characterized by severe symptoms (*52–54*). Since we found that eosinophils directly regulate neutrophils and GPR43 is a critical suppressor of eosinophil hyperactivation, our results offer new insights into mechanisms that may underlie the development of mixed-granulocytic asthma. For example, our findings suggest that insufficient GPR43 signaling due to antibiotic overuse or inadequate dietary fiber intake could induce dysfunctional eosinophils that aberrantly recruit and activate neutrophils in response to asthmogenic signals, which in turn trigger powerful Th17 responses. Further studies on the GPR43-eosinophil-Siglec-F^hi^ neutrophil-Th17 axis in not only asthma but also other diseases that involve both eosinophils and neutrophils (e.g. chronic obstructive pulmonary disease, infections, and tumors) are warranted.

## MATERIALS AND METHODS

### Experimental animals

*Gpr43*^fl/fl^ mice were generated using ES cells (Ffar2tm1a(EUCOMM)Hmgu) from EUCOMM. *Epx*-Cre mice (*55*) and Rosa26-eYFP^fl/fl^ mice (Jackson laboratory, 006148) were obtained from the late James J. Lee (Mayo Clinic) and Suk-jo Kang (KAIST), respectively. All mice were on the C57BL/6 background. Control and experimental groups were co-housed from 3 weeks after birth until the end of the experiments. Eight to 13-week old male mice were used in experiments. Mice were maintained under specific pathogen-free conditions in the KAIST Laboratory Animal Resource Center. All mouse experiments were approved by the KAIST Institutional Animal Care and Use Committee (IACUC).

### Allergic airway inflammation models

For the HDM-induced asthma model, 10 µl of 1 mg/ml house dust mite extract (Greer, USA) dissolved in PBS was administered intranasally to mice on days 0, 7, 8, 9, 10, and 11. In the case of the OVA-induced asthma model, mice were sensitized intraperitoneally on day 0 with 200 µl of 1 mg/ml OVA (Sigma-Aldrich, USA) and Alum (Imject, USA) mixed in a 1:1 ratio. For the challenge, 20 µl of 2.5 mg/ml OVA in PBS was administered intranasally on days 7, 8, and 9. For intranasal administration, mice were anesthetized by intraperitoneal injection of a mixture containing ketamine (80 mg/kg) and rompun (4 mg/kg) in PBS. In both models, mice were euthanized one day after the last challenge using CO_2_.

### Measurement of airway resistance

Airway resistance was measured using FlexiVent (SCIREQ, Canada). Mice were anesthetized by intraperitoneal injection of pentobarbital sodium solution in PBS (30 mg/kg) and intubated with 20-gauge catheters by tracheostomy. To block spontaneous breathing, 100 µl of 1 µg/ml pancuronium bromide (Sigma-Aldrich) was injected intraperitoneally. Mice were mechanically ventilated at a tidal volume of 10 ml/kg and a respiratory rate of 150/min. Respiratory resistance (Rrs) values were acquired after administering increasing doses (0, 5, 10, 20 mg/ml) of aerosolized methacholine (Sigma-Aldrich).

### Histological analysis of asthmatic lung

Half of the left lung tissues were excised, fixed with 3.7% formaldehyde (Sigma-Aldrich) at 4 °C overnight, and embedded into paraffin using a tissue processor (TP1020 Automatic Benchtop Tissue Processor / HistoCore Arcadia, Leica). The paraffin block was cut into 10 µm-thick sections with the microtome (RM2235, Leica) and stained with hematoxylin (Merck, Germany) and eosin (Merck) using a standardized protocol. Periodic Acid Schiff (PAS) and Picro Sirius Red staining were performed using kits (Abcam, UK) according to the manufacturer’s protocols. Images were obtained using a slide scanner (Pannoramic SCAN II, 3DHISTECH, Hungary) in the EM & histology core facility at the KAIST biomedical research center.

### Immunofluorescence imaging of asthmatic lung

After euthanizing the mice, 1 ml OCT solution (Sakura Finetek, USA) was injected into the lungs through the trachea, and the lungs were extracted. The extracted lungs were immersed in OCT solution and preserved at −20°C. OCT-embedded lungs were sectioned into 14 µm-thick slices and attached to the slide glasses. After fixation with 4% paraformaldehyde (Sigma-Aldrich) and permeablization with 0.3% Triton X-100 in PBS, anti-Siglec-F (E50-2440, BD Biosciences) and anti-myeloperoxidase (P11247, R&D Systems) antibodies were used to stain eosinophils and neutrophils, respectively. Anti-rat IgG-AF594 and anti-goat IgG-AF488 antibodies (Thermo Fisher Scientific) were used as the secondary antibodies. After nuclear staining with DAPI (Thermo Fisher Scientific) and adding a fluorescence mounting medium (Dako, Denmark), a cover glass was mounted. Confocal imaging was performed using Nikon A1 HD25 microscope (Japan) and images were processed with NIS-Elements BR 4.60. software.

### Lung and BALF cell preparation

Lungs were chopped into 1 mm^3^ fragments and digested with 400 MandlU/ml collagenase D (Roche, Switzerland) and 10 µg/ml DNase I (Roche) in RPMI1640 media (Welgene, South Korea) containing 3% FBS (Atlas biologicals, USA), 20 mM HEPES (Welgene), 1 mM non-essential amino acids (Welgene), 100 U/ml penicillin, 100 µg/ml streptomycin (Welgene), and 1 mM sodium pyruvate (Welgene) at 37°C for 30 min with stirring. The enzyme reaction was stopped by adding 10 µl of 0.5 M EDTA (Sigma-Aldrich) and incubating the cells for an additional 5 min. After vigorous resuspension, the cells were filtered through 100 µm strainers (SPL, South Korea). Red blood cells were lysed with ACK lysis buffer containing 0.15 M NH_4_Cl (Daejung, South Korea), 10 mM KHCO_3_ (Daejung), and 0.1 mM EDTA (Sigma-Aldrich) for 3 min at room temperature. BALF was collected by flushing the airways with 1 ml PBS. To evaluate the tissue localization of immune cells by flow cytometry, mice were injected intravenously with 1 µg of CD45-PE antibody (30-F11, Biolegend) 5 min before sacrifice.

### Flow cytometry

Cells were first stained with Ghost dye violet 510 (Tonbo, USA) for live/dead staining and then with fluorescent dye-labeled antibodies in PBS-based buffer supplemented with 3 % FBS, 1 mM EDTA, and 20 mM HEPES on ice for 20 min. For intracellular cytokine staining, the cells were stimulated with a cocktail containing 81 nM PMA (Sigma-Aldrich), 1.34 µM ionomycin (Sigma-Aldrich), 2 µM monensin (Sigma-Aldrich), and 10.6 µM brefeldin A (Sigma-Aldrich) at 37°C for 4 h. The cells were then fixed and permeabilized with 3.7% formaldehyde (Sigma-Aldrich) and 0.5% saponin (Sigma-Aldrich), respectively. To stain the transcription factors in CD4 T cells, a FoxP3 staining kit (eBioscience) was used according to the manufacturer’s protocol. The following antibodies were used. CD45.2-APC-Cy7 (104), CD4-APC-H7 (GK1.5), Siglec-F-AF647 (E50-2440), IL-10-APC (JES5-16E3), RORγt-AF647 (Q31-378), I-A/I-E--FITC (2G9), CD11c-FITC (HL3), CD11b-FITC (M1/70), PECAM-1-FITC (MEC13.3), IL-4-PE (11B11), Siglec-F-PE (E50-2440) (all from BD Biosciences), CD45.2-PB (104), CD45-PE (30-F11), Ly6c-PE (HK1.4), Ly6G-PerCP-Cy5.5 (1A8), T-bet-PE-Cy7 (4B10), TCRβ-FITC (H57-597), CD11c-BV785 (N418), PECAM-1-APC (MEC13.3) (all from Biolegend), MHCII-eFluor 450 (M5/114.15.2), CD11b-PB (M1/70), Gr-1-APC (RB6-8C5), Gr-1-PE-Cy7 (RB6-8C5), GATA3-PE (TWAJ), IL-17A-PerCP-Cy5.5 (eBio17B7), TCRβ-PE-Cy7 (H57-597), F4/80-PE-Cy7 (BM8), IL-13-eFluor660 (eBio13A), IL-5-PE (TRFK5), FoxP3-PerCP-Cy5.5 (FJK-16s) (all from eBioscience), and CD4-violetFluor 450 (GK1.5) (Tonbo). Flow cytometric analysis was performed using BD Biosciences LSR Fortessa X-20 and data were analyzed with BD Biosciences Flowjo v.10.10.0.

### Purification of lung eosinophils and morphological analysis

After staining lung cells with fluorescent dye-labeled antibodies, eosinophils were purified by fluorescence-activated cell sorting using BD Biosciences Aria Fusion. Dead cells were excluded using propidium iodide staining. The purity of the sorted eosinophils was over 95%. For morphological analysis, 50,000 cells were attached to slide glass using Cytospin 4 (Thermo Fisher Scientific). After overnight drying, cells were fixed with 3.7% formaldehyde, stained with hematoxylin and eosin (Merck) using a standardized protocol, and mounted with xylene mounting solution (BBC biochemical, USA). To analyze their nuclear shape, cells were stained with propidium iodide and mounted with a fluorescence mounting medium (Dako, Denmark). The images were taken with Pannoramic SCAN II.

### Purification of bone marrow and lung neutrophils

To purify bone marrow neutrophils, bone marrow cells were collected by flushing the tibia and femur with RPMI1640-based media containing 2% FBS (Atlas biologicals), 10 mM HEPES, and 100 U/ml penicillin, and 100 µg/ml streptomycin using 26 G syringes. The bone marrow and lung cell preparations were enriched for neutrophils by negative selection with biotin-conjugated B220 (RA3-6B2), CD3ε (145-2C11), Ter119 (Ly-76), NK1.1 (PK136) (all from Biolegend), CD19 (1D3, BD Biosciences), and TCRγδ (eBioGL3, eBioscience) antibodies by using EasySep Mouse Streptavidin RapidSpheres isolation kit (STEMCELL Technologies). The neutrophils were further stained with fluorescent dye-labeled antibodies and purified using Aria Fusion cell sorter (BD Biosciences). Their purity was over 95%.

### In vitro differentiation of Siglec-F^hi^ neutrophils

For cytokine-induced differentiation of Siglec-F^hi^ neutrophils, 0.2 million neutrophils purified from the bone marrow were cultured in RPMI-based media supplemented with 10% FBS (Hyclone), 1 mM non-essential amino acids, 5 mM HEPES, 100 U/ml Penicillin, 100 µg/ml Streptomycin, 1 mM sodium pyruvate, 50 µg/ml gentamicin (Gibco, USA), 55 µM β-mercaptoethanol (Sigma-Aldrich), and 2 mM L-glutamine (Gibco) for 2 days in the presence of 10 ng/ml of each cytokine, including IL-1α, IL-1β, IL-1Ra, IL-2, IL-4, IL-5, IL-8, IL-10, IL-12, IL-17A, IL-33, IFNγ (R&D Systems), IL-6, TNF (BD Biosciences), GM-CSF (Peprotech) M-CSF (Sigma-Aldrich), and TGFβ (Miltenyi Biotec), or 10 µg/ml LPS (InvivoGen).

For eosinophil-induced differentiation of Siglec-F^hi^ neutrophils, 0.2 million bone marrow neutrophils were co-cultured with 0.1 million eosinophils that were purified either from the steady-state or asthmatic lungs in the culture media described above in 96-well round bottom plate for 2 days. For antibody-mediated blocking of Siglec-F^hi^ neutrophil differentiation, eosinophils were pre-treated with 5 µg/ml anti-IL-4 (BVD4-1D1, BD Biosciences) and 5 µg/ml anti-GM-CSF (MP122E9, R&D Systems) or 1 µg/ml anti-PECAM-1 (MEC13.3, Biolegend) antibodies for 5 min and co-cultured with neutrophils.

### Analysis of eosinophil-neutrophil interaction

Bone marrow neutrophils (0.2 million) were cultured in the presence or absence of asthmatic lung eosinophils (0.1 million) in a 96-well flat bottom plate for 2 days, and cells were imaged using a Nikon ECLIPSE Ts2R microscope (Japan). For fluorescence imaging, bone marrow neutrophils labeled with carboxyfluorescein succinimidyl ester (Thermo Fisher Scientific) and asthmatic lung eosinophils (0.1 million each) were co-cultured in a 96-well round bottom plate for 4 h. Without any resuspension, cells were carefully transferred into the cytofunnel and attached to a slide glass using Cytospin 4 (Thermo Fisher Scientific). After overnight drying, the cells were fixed with 4% paraformaldehyde (Sigma-Aldrich), permeabilized with 0.3% Triton X-100 in PBS, and stained with an anti-CD63-APC antibody (NVG-2, Biolegend). Fluorescence images were taken using a Nikon A1 HD25 confocal microscope.

### Neutrophil transwell migration assay

Bone marrow neutrophils were labeled with 5 µM CellTrace Violet (CTV, Invitrogen) at 37°C for 20 min. CTV-labeled neutrophils (0.1 or 0.2 million) and asthmatic lung eosinophils (0.7 million) were placed in a 3 µm transwell insert (Corning) and the bottom 96-well plate, respectively, and cultured for 24 h. The cells in the bottom plate were resuspended to dissociate the aggregated cells, and CTV-labeled neutrophils were imaged using a Nikon ECLIPSE Ts2R fluorescence microscope.

### In vitro Th17 differentiation

To determine the ability of neutrophils to induce Th17 differentiation, Siglec-F^hi^ and Siglec-F^low^ neutrophils were prepared by culturing bone marrow neutrophils for 2 days in the presence or absence of 10 ng/ml GM-CSF (Peprotech) and 10 ng/ml IL-4 (R&D Systems), respectively. To prepare naïve CD4 T cells, spleens and peripheral lymph nodes were ground through 100 µm strainers using the piston of 1 ml syringes. After lysing red blood cells with ACK lysis buffer, naïve CD4 T cells were isolated by negative selection with biotin-conjugated CD8a (53-6.7), CD11b (M1/70), B220 (RA3-6B2), Ter119 (Ly-76), CD24 (M1/69), CD25 (PC61), CD44 (IM7) (all from Biolegend), TCRγδ (eBioGL3, eBioscience), CD49b (DX5, eBioscience), CD19 (1D3, BD Biosciences), and CD11c (HL3, BD Biosciences) antibodies using EasySep Mouse Streptavidin RapidSpheres isolation kit (STEMCELL Technologies). Naïve CD4 T cells (0.1million) were co-cultured with Siglec-F^hi^ or Siglec-F^low^ neutrophils (0.05 million) in the presence of 5 ng/ml TGFβ (Miltenyi Biotec), 20 ng/ml IL-6 (BD Biosciences), 5 µg/ml anti-IL-4 (BVD4-1D1, BD Biosciences), 2.5 µg/ml anti-IL-2 (S4B6, BD Biosciences), 5 µg/ml anti-IFNγ (R4-6A2, Invitrogen) in a 96-well flat bottom plate pre-coated with 2.5 µg/ml anti-CD3 (145-2C11, BD Biosciences) and 1 µg/ml anti-CD28 (37.51, BD Biosciences). After 3 days, the frequencies of RORγt or IL-17A-expressing CD4 T cells were analyzed by flow cytometry.

To determine the ability of eosinophils to induce Th17 differentiation, naïve CD4 T cells (0.1 million) were cultured as described above except the neutrophils were substituted with asthmatic lung eosinophils (0.1 million). To assess whether eosinophils indirectly influence Th17 differentiation *via* dendritic cells, splenic dendritic cells were added to the naïve CD4 T cell-eosinophil co-culture. To prepare the splenic dendritic cells, spleens were chopped into 1 mm^3^ fragments and digested with 400 MandlU/ml Collagenase D and 10 µg/ml DNase I in RPMI1640 media containing 3% FBS (Atlas biologicals), 20 mM HEPES, 1 mM non-essential amino acids, 100 U/ml penicillin, 100 µg/ml streptomycin, and 1 mM sodium pyruvate at 37°C for 30 min with stirring. The cells were centrifuged over 17.5% accudenz solution at 805 g for 20 min at room temperature, and the cells at the interface were collected. The dendritic cells were further stained with fluorescent dye-labeled antibodies and purified using BD Biosciences Aria Fusion cell sorter to a purity of over 95%.

### Cytokine ELISA

To measure cytokine production from T cells in the lung, total asthmatic lung cells (0.5 million) were cultured in flat bottom 96-well plates coated with 1 µg/ml of anti-CD3 (145-2C11, BD Biosciences) and 1 µg/ml anti-CD28 (37.51, BD Biosciences) for 3 days, and the culture supernatants were collected. To measure neutrophil cytokine production, bone marrow neutrophils (0.3 million) were cultured in the presence of 10 ng/ml IL-4 (R&D Systems) or 10 ng/ml GM-CSF (Peprotech) in 96-well round bottom plates for 2 days, and the culture supernatants were collected. On the second day of the culture, neutrophils were either stimulated or not with 100 nM PMA (Sigma-Aldrich) and 1 µg/ml ionomycin (Sigma-Aldrich). The cytokine ELISAs involved the following capture and detection antibodies. IL-4: anti-mouse IL-4 (11B11, BD Biosciences), biotinylated anti-mouse IL-4 (BVD6-24G2, BD Biosciences); IL-5: anti-mouse IL-5 (TRFK5, BD Biosciences), biotinylated anti-mouse IL-5 (TRFK4, BD Biosciences); IL-13: anti-mouse IL-13 (38213, R&D Systems), biotinylated anti-mouse IL-13 (P20109, R&D Systems); IL-17A: anti-mouse IL-17A (TC11-18H10, BD Biosciences), biotinylated anti-mouse IL-17A (TC11-8H4, BD Biosciences); IFNγ: anti-mouse IFNγ (AN-18, Invitrogen), biotinylated anti-mouse IFNγ (R4-6A2, Invitrogen); TNF: anti-mouse TNF (TN3-19.12, BD Biosciences), biotinylated anti-mouse TNF (516D1A1, BD Biosciences); and IL-1α: anti-mouse IL-1α (ALF-161, Biolegend), biotinylated anti-mouse IL-1α (BL1a-89, Biolegend) antibodies. IL-10 was measured using an IL-10 ELISA kit (eBioscience). Half-Area 96-well microplates (Corning) were coated with 2 µg/ml capture antibodies in PBS at 4°C overnight and blocked with blocking buffer containing 2% BSA fraction V (Roche) in PBS-T (0.05% Tween-20 (Daejung) in PBS) for 1 h at room temperature. Subsequently, samples were added and incubated for 1 h. Thereafter, 1 µg/ml biotin-conjugated antibodies were added for 45 min, followed by treatment with 50 ng/ml streptavidin-horseradish peroxidase (R&D Systems) for 20 min. Between each step, plates were washed at least 3 times with PBS-T. To develop the reaction, 50 µl of tetramethylbenzidine solution (Surmodics, USA) was added, and the reaction was stopped by adding 50 µl of 0.5 M H_2_SO_4_ (Daejung). Absorbance at O.D.450 nm was read using SpectraMax M2 Microplate Reader (Molecular Devices, USA).

### Preparation of bone marrow-derived eosinophils

Bone marrow-derived eosinophils (BMDEs) were differentiated by following the previously described protocol(*56*). In brief, bone marrow cells (3 x 10^6^/ml) were differentiated with 100 ng/ml SCF (Peprotech), and 100 ng/ml FLT3 ligand (R&D Systems) in RPMI1640 media containing 20 % FBS (Hyclone), 1 mM non-essential amino acids, 25 mM HEPES, 100 U/ml Penicillin, 100 µg/ml streptomycin, 1 mM sodium pyruvate, 50 µM β-mercaptoethanol, and 2 mM L-glutamine in 6-well plates for 4 days. The mediium was then changed to new culture medium containing 10 ng/ml IL-5 (R&D Systems). On days 8 and 11, the medium was changed again with new IL-5-containing medium, and the cell number was adjusted to 1 x 10^6^/ml. On day 14, Ca^2+^ flux assay and RNA extraction were performed.

### Ca^2+^ flux assay

Cells were loaded with 2 µg/ml Fluo-3 AM (Thermo Fisher Scientific) and 5 µg/ml Fura Red AM (Thermo Fisher Scientific) in RPMI1640 medium containing 2% FBS, 25 mM HEPES, 2.5 mM probenecid (Sigma-Aldrich), and 0.02% pluronic F-127 (Sigma-Aldrich) at 37°C for 20 min. After washing, the cells were analyzed using a flow cytometer (LSR Fortessa X-20, BD Biosciences). Basal fluorescence was recorded for 30 seconds and changes in the fluorescence were recorded for an additional 3 min after stimulating the cells with 10 mM acetate.

### RNA extraction

Asthmatic lung eosinophils, BMDEs, and bone marrow neutrophils were lysed in Qiazol (QIAzen, Germany), and chloroform (Daejung) was added in a 5:1 ratio. After centrifugation, the clear upper phase was collected and RNA was precipitated by adding the same volume of 100% isopropanol (Merck) and 20 µg glycogen (Sigma-Aldrich). After centrifugation, the pellet was washed with 75% ethyl alcohol (Merck) diluted in DEPC water (Biosesang, South Korea) and air-dried. RNA was solubilized in nuclease-free water (Invitrogen).

### Quantitative PCR

RNAs were reverse-transcribed to cDNA using GoScript™ reverse transcriptase, ribonuclease inhibitor, dNTP, MgCl2, and oligo(dT)15 primer (all from Promega). Using cDNAs as templates, qPCR analysis of genes of interest was performed with SYBR Green Realtime PCR Master Mix (TOYOBO, Japan) and CFX Connect Real-Time System (Bio-Rad). Gene expression levels were calculated based on the comparative Cq method and normalized to *Hprt* gene expression. Primers are described in Table S2.

### Bulk-RNA sequencing

The integrity of asthmatic lung eosinophil RNA was measured using a Bioanalyzer 2100 (Agilent Technologies, USA). Ribosomal RNAs were removed using NEBNext rRNA Depletion Kit v2 (USA), and the cDNA library was synthesized using NEBNext Ultra II Directional RNA Library Prep kit according to the manufacturer’s protocols. Sequencing was conducted on an Illumina HiSeq2500 with paired-ended reads of 100 base pairs by Macrogen (South Korea). Fastq files were trimmed with package Fastp and aligned to the mm10 mouse reference genome sequences with STAR. Transcript per million (TPM) was calculated from the counts obtained with FeatureCounts and differentially expressed genes (DEGs) were acquired from the raw counts with DESeq2 by using a p-value below 0.05. All processes were conducted through Galaxy (https://usegalaxy.org/). Additional pathway analysis was performed using pathfindR and Clusterprofiler in R v.4.3.1. Publicly available bulk-RNA sequencing data for eosinophils from mouse spleen(*25*), peritoneal cavity(*25*), lung(*23*), colon(*24*), skin(*14*), blood(*14*), bone marrow(*14*), BMDE(*26*), and human peripheral blood(*27*) were analyzed with the same processes.

### Single-cell RNA sequencing

Asthmatic lung cells from *Epx*^Cre/+^ and *Epx*^Cre/+^*Gpr43*^fl/fl^ mice were isolated from 3 mice per genotype, pooled, and stained with Ghost dye e780. Live cells, excluding FSC^hi^ SSC^hi^ cells, were sorted and suspended in PBS containing 0.04% BSA. The total of 28,320 and 25,620 cells from *Epx*^Cre/+^ and *Epx*^Cre/+^*Gpr43*^fl/fl^ mice, respectively, were loaded onto Chromium-controller (10X Genomics), and scRNA-seq libraries were generated by using the Chromium Next GEM Single Cell 3ʹ Reagent Kits v 3.1 (10X Genomics) according to the manufacturer’s protocol at the KAIST NGS core facility. Sequencing was performed on a HiSeqX platform by Macrogen (South Korea).

Raw sequencing files were initially processed with the Cell Ranger (v.4.0.0) pipeline, using the mm10 reference, and downstream analysis was conducted based on the Scanpy (v.1.8.2) pipeline. Low-quality cells meeting any one of the following criteria were excluded in the downstream analysis: (1) UMI gene count < 1000, (2) number of genes detected < 500, (3) number of genes detected > 7000, (4) fraction of mitochondrial genes > 10%, and (5) predicted doublets identified with the Scrublet (v.0.2.2) package. Raw counts were normalized in each cell (100,000 counts per cell) and log-transformed (log1p). Highly variable genes (n=2,418) were selected by using the highly_variable_genes function in the Scanpy package. Principal component analysis (PCA) was conducted (n_PCs = 50) on the scaled expression values of the highly variable genes. Neighborhood graphs of the cells were constructed (n_neighbors = 15) based on PC coordinates, and the Uniform Manifold Approximation and Projection (UMAP) embedding of the cells was calculated. After the preprocessing steps, 3,989 cells from the *Epx*^Cre/+^ sample and 4,871 cells from the *Epx*^Cre/+^*Gpr43*^fl/fl^ sample were retrieved and analyzed. Wilcoxon rank-sum test was used for the DEG analysis, and p-values were adjusted with the Benjamini-Hochberg method.

For gene set enrichment analysis (GSEA) of DEGs between Siglec-F^low^ and Siglec-F^hi^ neutrophils, a pre-ranked gene set was created by listing 3515 DEGs with an adjusted p-value of less than 0.05, ranked by test statistics score using GSEApy v.1.1.3. Normalized enrichment scores (NES) and false discovery rates (FDR) were obtained by using following parameters: (1) number of permutations = 1000, (2) permutation type: gene set, and (3) weighted enrichment statistics. Significant gene sets were those with an FDR < 0.25 and p-value < 0.05. Gene sets for the analysis were obtained from the Kyoto Encyclopedia of Genes and Genomes (KEGG) and Gene Ontology (GO) databases. To compare the Siglec-F^hi^ neutrophils identified in our HDM-induced asthma model to those in other disease models using publicly available transcriptomic data(*31–33*), another pre-ranked gene set was generated by ranking the same 3515 DEGs according to LogFC values. Gene lists for the Siglec-F^hi^ neutrophil signature genes in the other disease models were generated, using the top 350 DEGs between Siglec-F^low^ and Siglec-F^hi^ neutrophils in each disease model, and ranked based on LogFC values.

Analysis of publicly available scRNA-seq data was performed using Seurat v.4.1.3. Mtx files for esophagus and duodenum samples of 10 eosinophilic esophagitis patients were downloaded (GSE175930)(*57*) together with feature and cell information files, and Seuratobject was created. Low-quality cells meeting any one of the following criteria were excluded in the downstream analysis: (1) min.cells < 3, (2) min.features < 200, (3) number of genes detected < 200, (4) number of genes detected > 6000, and (5) fraction of mitochondrial genes > 20%. Seuratobjects were normalized, and highly variable genes were found by using the vst selection method. Each esophagus and duodenum sample was integrated separately (n_dimension = 50), and PCA was conducted (n_PC = 30). Neighborhood graphs of the cells were constructed (n_dimenstion = 30), UMAP was created, and the cells were clustered (esophagus: resolution = 0.3; duodenum: resolution = 0.1). Each cluster was manually annotated using known marker genes.

### Statistical analysis

Statistical analyses were performed with GraphPad Prism v.8.4.2.

## Acknowledgments

The authors thank the scientific contributions and technical assistance by Dr. Yongsuk Hur at the EM & Histology Core Facility and Ms. Jiye Kim at NGS Core Facility in the BioMedical Research Center, KAIST.

## Author contributions

YMK and JY conceived the project and wrote the manuscript with comments from all authors. YMK supervised the study. JY performed overall experiments and sequencing data analysis. SK performed single-cell RNA sequencing data analysis. HSS performed confocal microscopy. HYK provided technical advices on *in vivo* experiments and critical comments on data interpretation.

## Data and materials availability

The raw data and processed files generated from bulk RNA sequencing of asthmatic lung eosinophils have been deposited in the Gene Expression Omnibus with accession code GSE271832. The raw data and processed files generated from single-cell RNA sequencing of whole asthmatic lung cells are available under accession code GSE271835. Source data are provided with this paper.

## Supplementary Materials

**Fig. S1.**
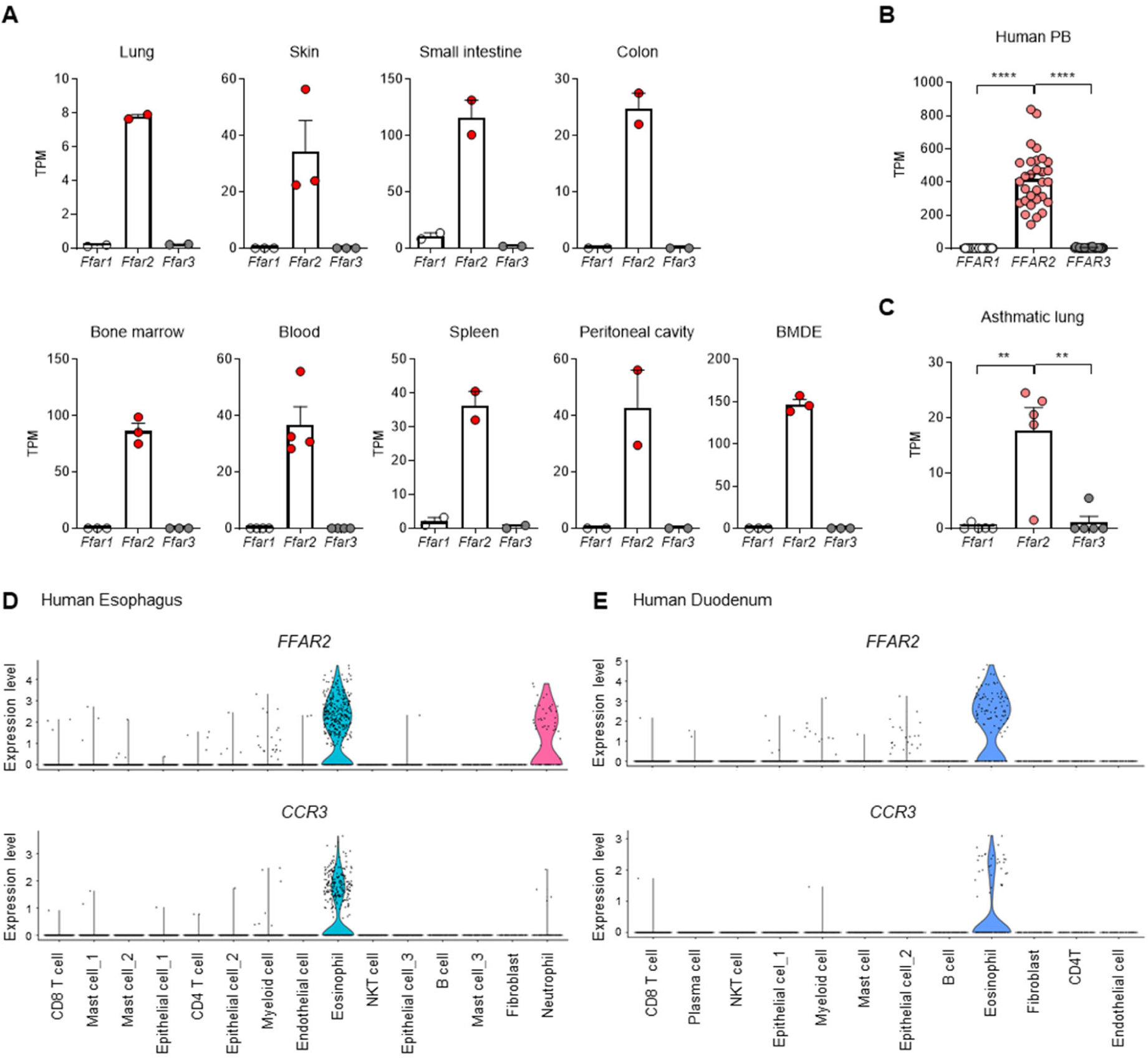
The expression of GPR43 in eosinophils. **A-C**, The expression of free fatty acid receptors encoding GPR40 (*Ffar1*/*FFAR1*), GPR43 (*Ffar2*/*FFAR2*), GPR41 (*Ffar3*/*FFAR3*) in eosinophils from various mouse organs and bone marrow-derived eosinophils (BMDE). lung: n = 2, skin: n = 3, small intestine: n = 2, colon: n = 2, bone marrow: n = 3, blood: n = 4, spleen: n = 2, peritoneal cavity: n = 2, BMDE: n = 3 (**A**), human peripheral blood (n = 30) (**B**), and mouse asthmatic lung (n = 5) (**C**). **D-E**, Violin plots for the expression of *FFAR2* and *CCR3* in each cell type of the esophagus (**D**) and duodenum (**E**) from eosinophilic esophagitis patients. Data are presented as mean ± s.e.m. Data in (A, B, and D) were analyzed using publicly available sequencing datasets

**Fig. S2.**
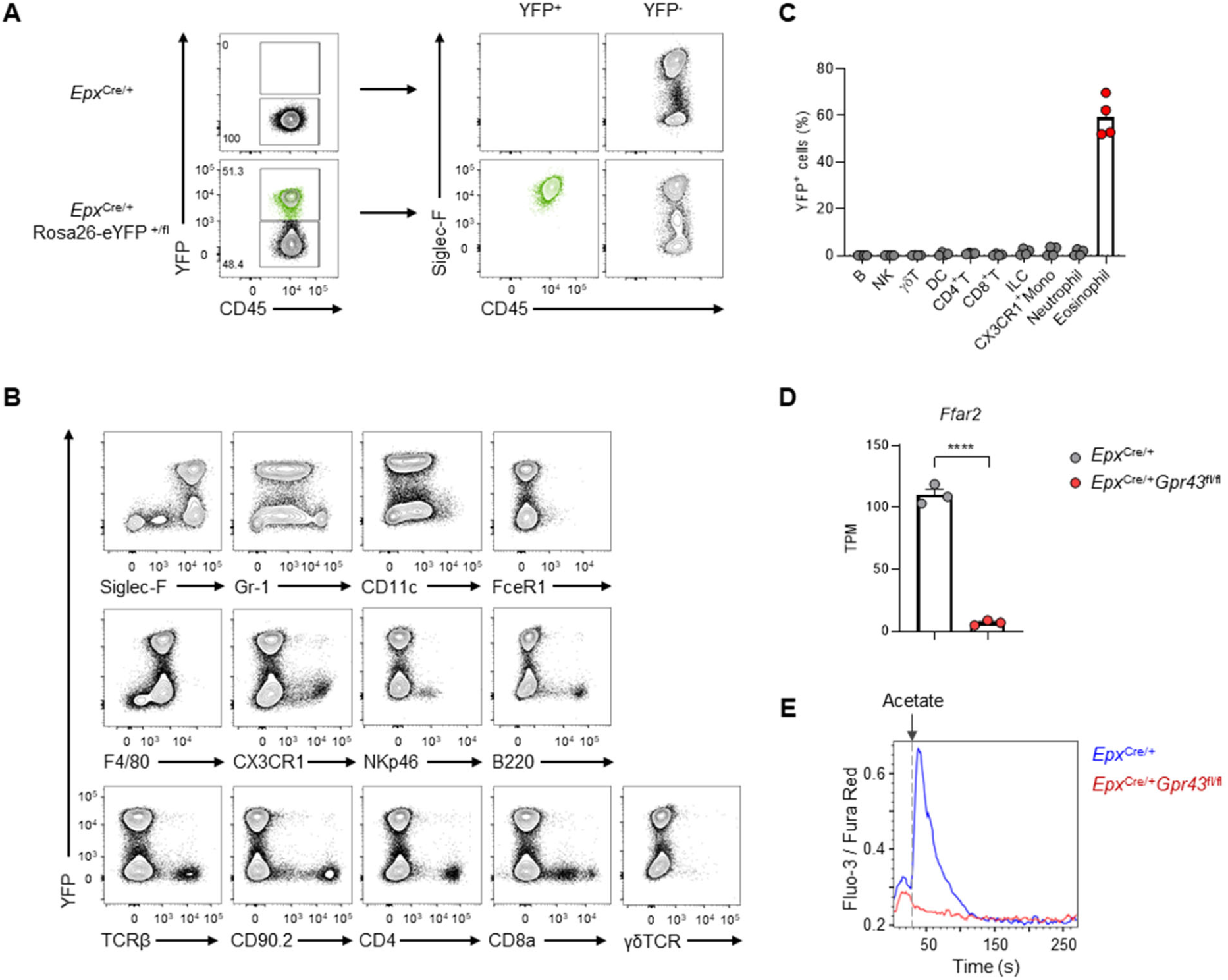
*Gpr43* is efficiently deleted in eosinophils of *Epx*^Cre/+^*Gpr43*^fl/fl^ mice. **A**, **B**, Representative plots showing YFP expression in eosinophils (**A**) and various surface marker-expressing immune cells (**B**) from asthmatic lungs of *Epx*^Cre/+^Rosa26-eYFP^+/fl^ mice. (**C**) The percentage of YFP^+^ cells in various immune cells from asthmatic lungs (n = 4). Data in A-C are representative of two independent experiments. (**D**) The expression of *Ffar2* in BMDEs from *Epx*^Cre/+^ (n = 3) and *Epx*^Cre/+^*Gpr43*^fl/fl^ (n = 3) mice. (**E)** Ca^2+^ influx triggered by acetate in BMDE. Data in (E) are representative of three independent experiments. Data are presented as mean ± s.e.m. Unpaired two-tailed Student’s t-test was performed for statistical analysis (**** p < 0.0001).

**Fig. S3.**
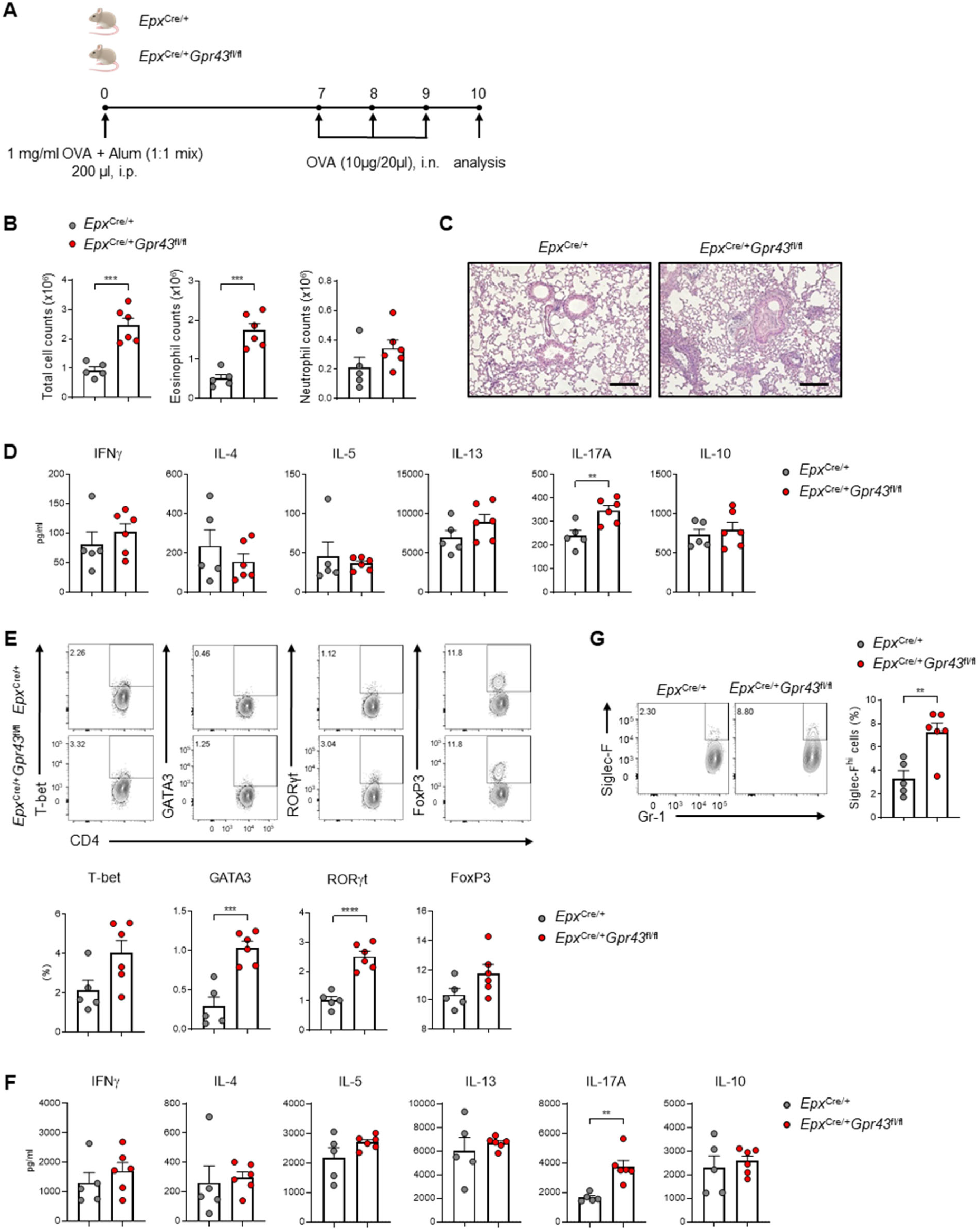
GPR43 deficiency in eosinophils causes more severe OVA-induced asthma. (**A**) Experimental scheme of OVA-induced asthma. (**B**) Total immune cell, eosinophil, and neutrophil count in BALF. (**C**) Histological analysis of immune cell infiltration (H&E) in the asthmatic lungs. Scale bar: 200 µm. (**D**) Cytokine levels in BALF. (**E**) The percentage of CD4 T cells expressing each transcription factor in asthmatic lungs. (**F**) Cytokine levels in the culture supernatant of asthmatic lung cells stimulated with anti-CD3 and anti-CD28. (**G**) The percentage of Siglec-F^hi^ cells among neutrophils in asthmatic lungs. *Epx*^Cre/+^ (n = 5), *Epx*^Cre/+^*Gpr43*^fl/fl^ (n = 6). Data are representatives of three independent experiments (A-G). Data are presented as mean ± s.e.m. Unpaired two-tailed Student’s t-test was performed for statistical analysis (** p < 0.01, *** p < 0.001, **** p < 0.0001).

**Fig. S4.**
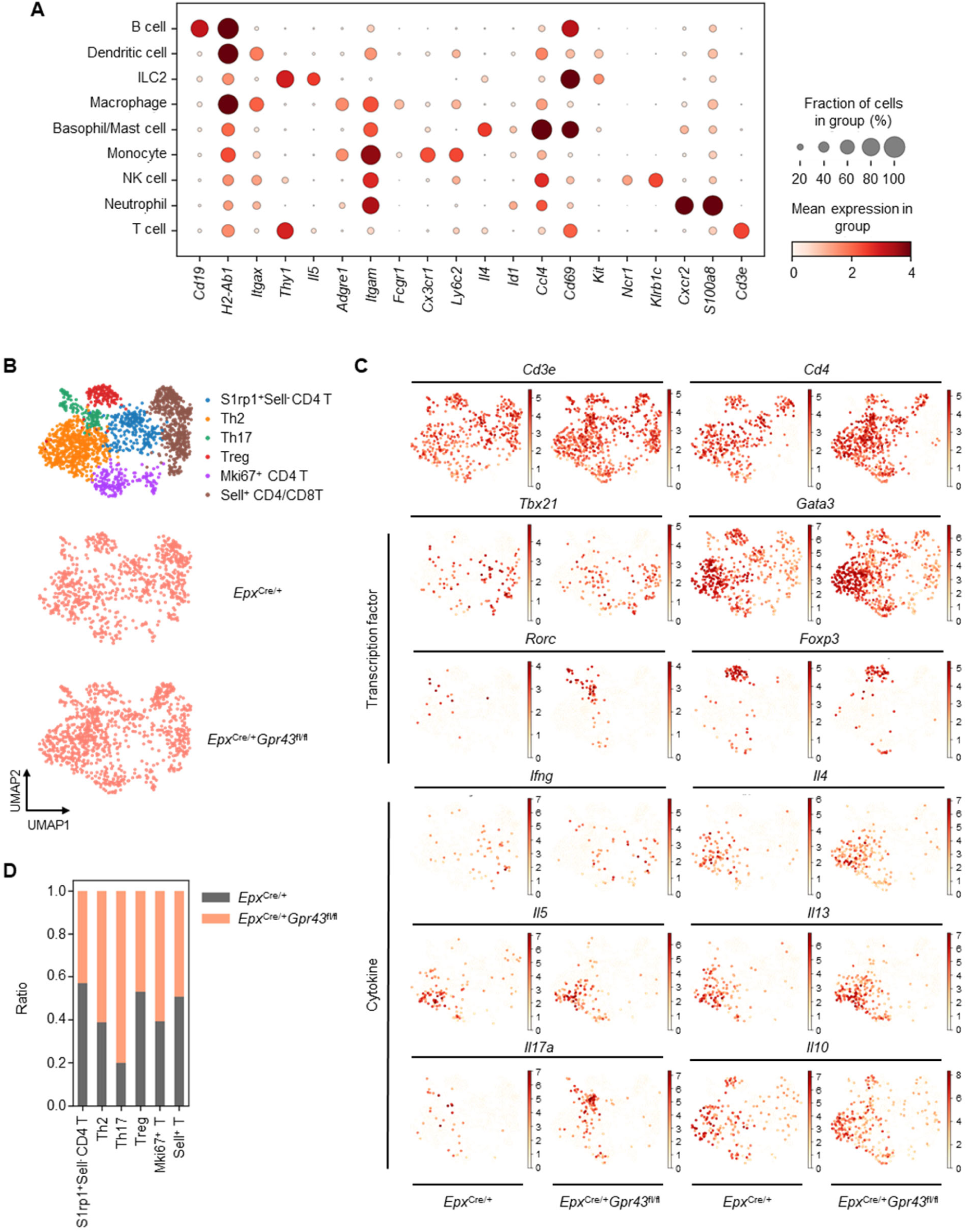
Increase of Th2 and Th17 cells in the asthmatic lungs of *Epx*^Cre/+^*Gpr43*^fl/fl^ mice. (**A**) Dot plot for representative marker genes of each cell cluster in the asthmatic lung. (**B**) UMAP plots presenting each T cell cluster in the asthmatic lungs. (**C**) UMAP plots displaying the marker gene expression of CD4 T subsets. (**D**) Ratio of T cells from *Epx*^Cre/+^ and *Epx*^Cre/+^*Gpr43*^fl/fl^ mice within each T cell cluster.

**Fig. S5.**
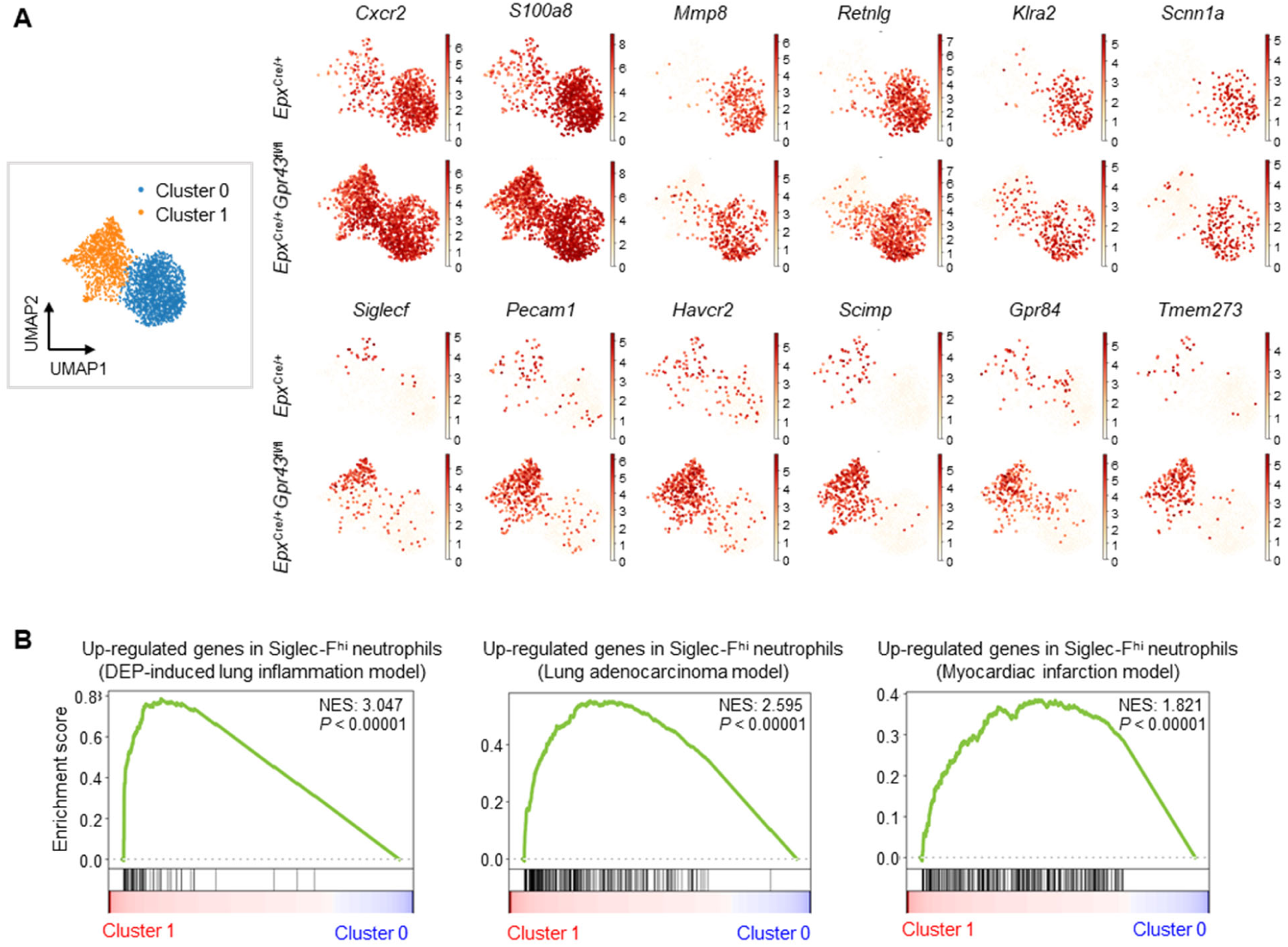
The analysis of differentially expressed genes in Siglec-F^hi^ and Siglec-F^low^ neutrophils. (**A**) UMAP plots presenting signature gene expression of Cluster 0 (Siglec-F^low^) and Cluster 1 (Siglec-F^hi^) neutrophils in asthmatic lungs. (**B**) Gene set enrichment analysis of DEGs between Siglec-F^hi^ and Siglec-F^low^ lung neutrophils from HDM-induced asthma model using gene sets each prepared with highly expressed genes in Siglec-F^hi^ neutrophils from three independent disease models.

**Fig. S6.**
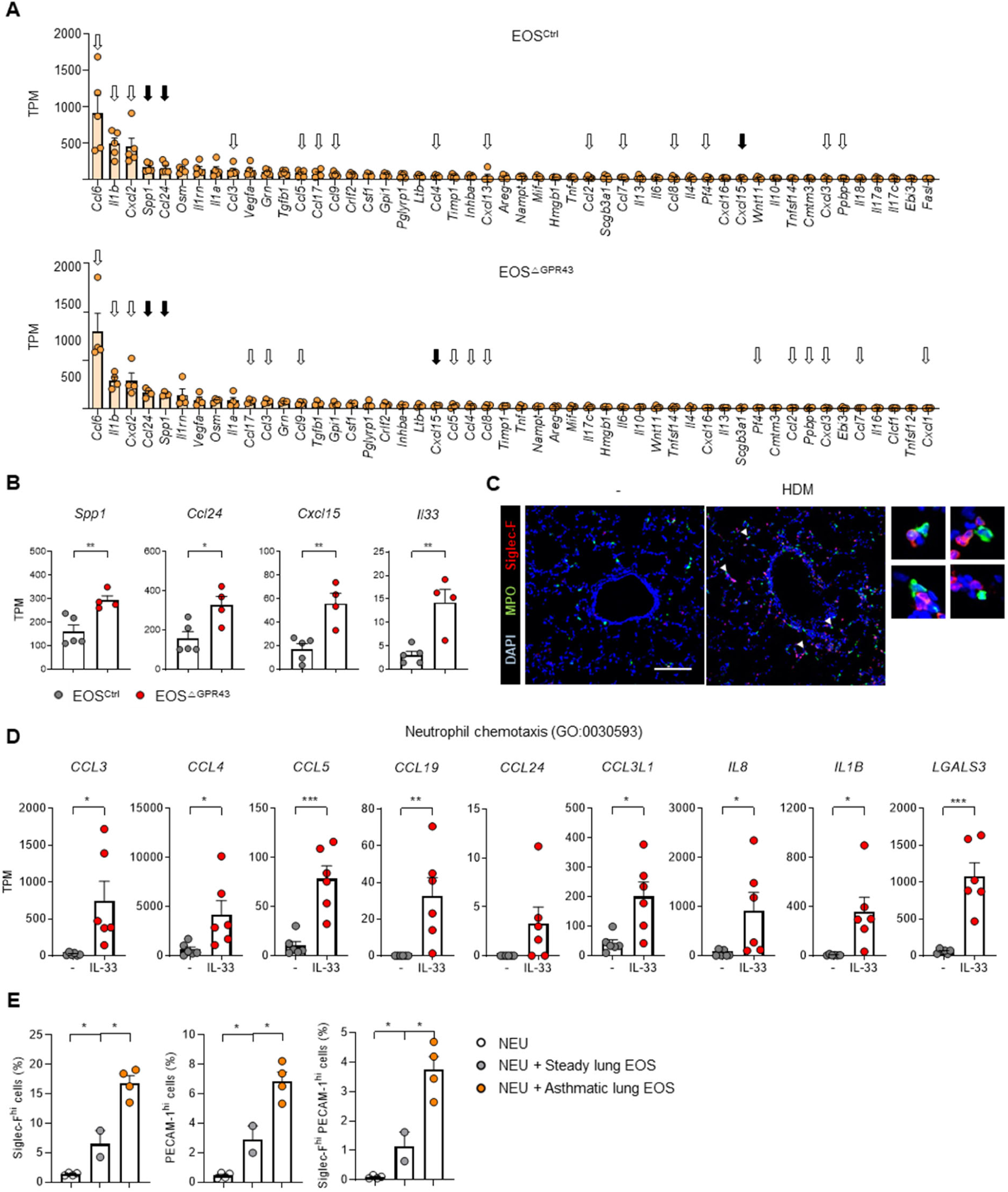
Eosinophils produce neutrophil chemoattractants. (**A**) The top 50 highly expressed cytokines and chemokines in asthmatic lung eosinophils. EOS^Ctrl^ (n = 5), EOS^△GPR43^ (n = 4). Genes related to neutrophil chemotaxis (GO_0030593) were marked with arrows, and black arrows indicate the ones increased in EOS^△GPR43^ compared to EOS^Ctrl^. (**B**) The expression of neutrophil chemoattractant genes in asthmatic lung eosinophils. EOS^Ctrl^ (n = 5), EOS^△GPR43^ (n = 4). (**C**) Fluorescence images of eosinophils (Siglec-F: red) and neutrophils (MPO: green) in the steady-state and asthmatic lungs. Interaction between eosinophils and neutrophils was marked by white arrowheads. Scale bar:100 µm. Data are representatives of two independent experiments. (**D**) The expression of genes related to neutrophil chemotaxis (GO_0030593) in human blood eosinophils treated with IL-33 (n = 6) or not (n = 6). Data was analyzed using publicly available RNA sequencing data. (**E**), The percentage of Siglec-F^hi^ and Siglec-F^hi^PECAM-1^hi^ cells among neutrophils co-cultured with either steady-state or asthmatic lung eosinophils. Steady-state (n = 2, each was pooled from 4 mice), asthma (n = 4). Data are presented as mean ± s.e.m. Unpaired two-tailed Student’s t-test was performed for statistical analysis (* p < 0.05, ** p < 0.01, *** p < 0.001).

**Fig. S7.**
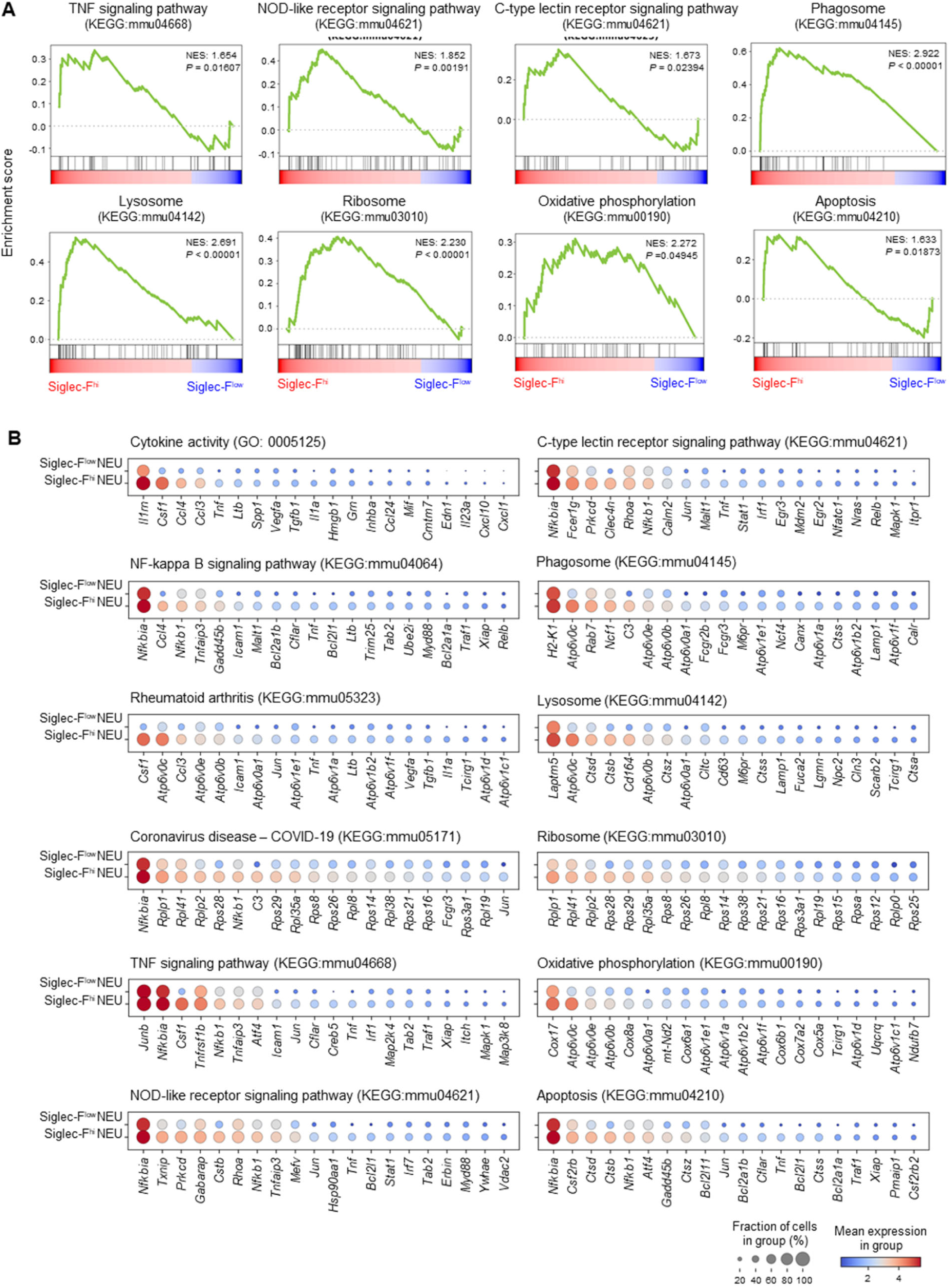
Molecular pathways enriched in Siglec-F^hi^ neutrophils. (**A**) Gene set enrichment analysis of DEGs between Siglec-F^low^ and Siglec-F^hi^ neutrophils. (**B**) Dot plots showing DEGs between Siglec-F^low^ and Siglec-F^hi^ neutrophils in each pathway.

**Fig. S8.**
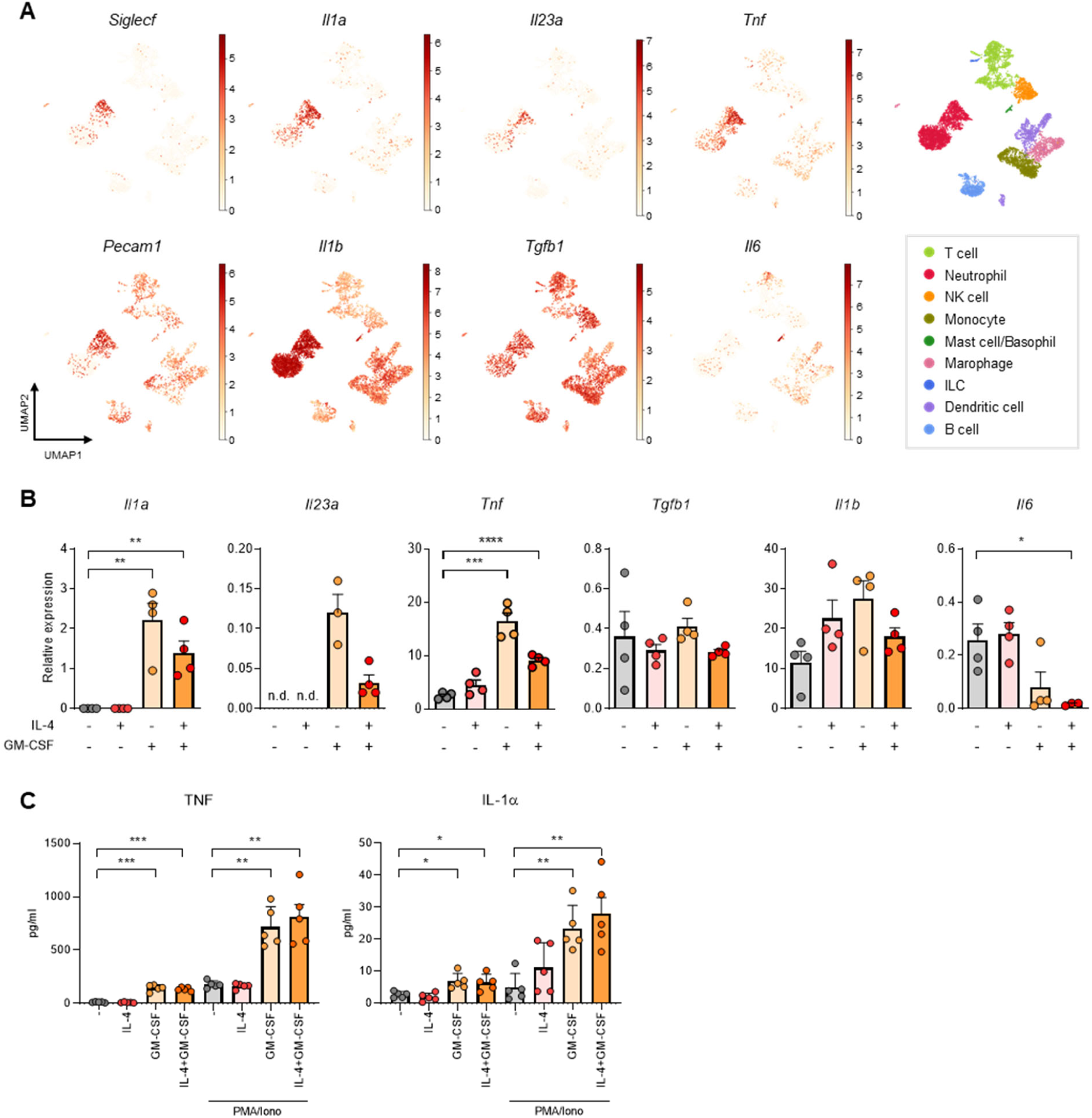
Siglec-F^hi^ neutrophils highly produce cytokines related to the differentiation of Th17 cells. (**A**) UMAP plots for genes related to Th17 differentiation. (**B**) The expression of genes related to Th17 differentiation in bone marrow neutrophils stimulated with IL-4 or GM-CSF (n = 3 or 4). (**C**) The levels of cytokines secreted from bone marrow neutrophils stimulated with IL-4 or GM-CSF (n = 5). Data are representative of two independent experiments. Samples in (B) are technical replicates, and samples in (C) are biological replicates. Data are presented as mean ± s.e.m. Unpaired two-tailed Student’s t-test (B) and paired two-tailed Student’s t-test (C) were performed for statistical analysis. (* p < 0.05, ** p < 0.01, *** p < 0.001, **** p < 0.0001).

**Fig. S9.**
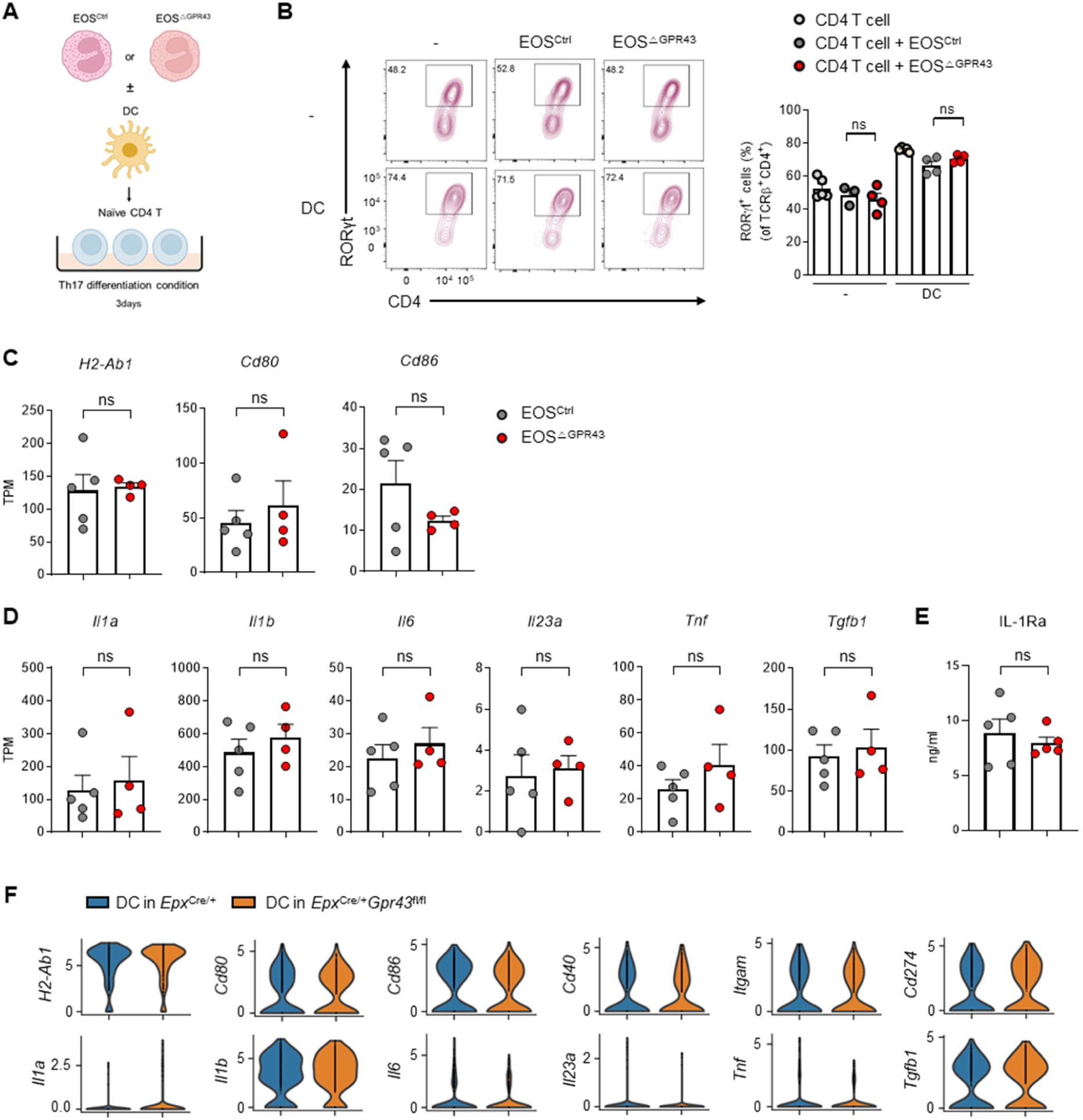
Eosinophils do not promote Th17 differentiation. (**A**) Experimental scheme for co-culture of asthmatic lung eosinophils and naïve CD4 T cells in the presence or absence of splenic DC under Th17 differentiation condition. (**B**) The proportion of RORγt^+^ cells after the co-culture. EOS^Ctrl^ (n = 3 or 4), EOS^△GPR43^ (n = 4). Data are representative of three independent experiments. **C**, **D**, The expression of genes related to antigen presentation (**C**) and Th17 differentiation (**D**) in asthmatic lung eosinophils. EOS^Ctrl^ (n = 5), EOS^△GPR43^ (n = 5). (**E**) IL-1Ra levels in the culture supernatant of asthmatic lung eosinophils. EOS^Ctrl^ (n = 5), EOS^△GPR43^ (n = 5). Data are representative of two independent experiments. (**F**) Violin plots for genes related to antigen presentation, activation, and Th17 differentiation in asthmatic lung dendritic cells. Data are presented as mean ± s.e.m. Unpaired two-tailed Student’s t-test was performed for statistical analysis (ns > 0.05).

**Fig. S10.**
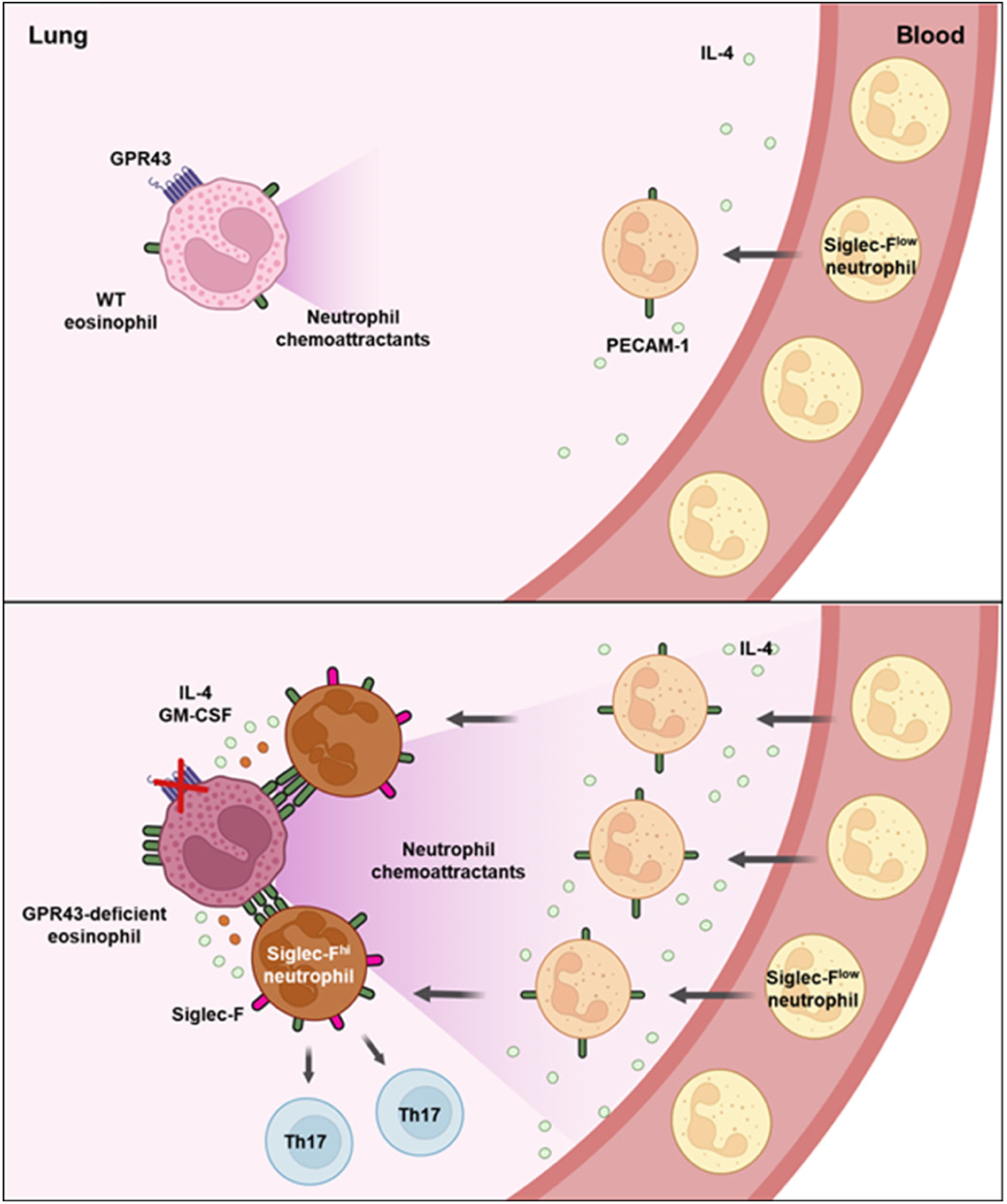
Schematics of eosinophil-mediated Siglec-F^hi^ neutrophil induction in asthmatic lungs. (Top) In wild-type mice, GPR43 suppresses eosinophil activation and the production of neutrophil chemoattractants, resulting in limited neutrophil recruitment and minimal Siglec-F^hi^ neutrophil generation. (Bottom) In eosinophil-specific GPR43-deficient mice, neutrophils infiltrating the lungs are exposed to higher levels of IL-4 and upregulate PECAM-1 expression. GPR43-deficient eosinophils are hyperactivated, produce more neutrophil chemoattractants, and express higher levels of PECAM-1 compared to wild-type eosinophils. This leads to increased neutrophil recruitment and stronger interactions between eosinophils and neutrophils, promoting the induction of Siglec-F^hi^ neutrophils via eosinophil-derived IL-4 and GM-CSF. Furthermore, Siglec-F^hi^ neutrophils enhance Th17 cell differentiation.

**Table S1.** Public dataset information.

| Sample | Sequencing Type | PMID | Accession number |
| --- | --- | --- | --- |
| Human blood eosinophils (Healthy and Asthmatics) | Bulk | 34475229 | E-MTAB-10190 |
| Asthmatics blood eosinophils (none vs IL-33 stimulation) | Bulk |  | E-MTAB-10189 |
| Mouse eosinophil Spleen | Bulk | 30686579 | SRR10060964<br>SRR10060965 |
| Mouse eosinophil Peritoneal cavity | Bulk |  | SRR10060962<br>SRR10060963 |
| Mouse eosinophil Lung | Bulk | 24837103 | SRR1206333<br>SRR1206334 |
| Mouse eosinophil colon | Bulk | 29758095 | SRR6233848<br>SRR6233851 |
| Mouse eosinophil BM-derived | Bulk | 32970801 | SRR11186563<br>SRR11186564<br>SRR11186565 |
| Mouse eosinophil Skin | Bulk | 37708282 | SRR21023657<br>SRR21023658<br>SRR21023659 |
| Mouse eosinophil Blood | Bulk |  | SRR21023667<br>SRR21023668<br>SRR21023669<br>SRR21023670 |
| Mouse eosinophil Bone marrow | Bulk |  | SRR21023671<br>SRR21023672<br>SRR21023673 |
| Human eophageal cells<br>Human duodenal cells (EoE patient) | Single cell | 34389613 | GSE175930 |
| Siglec-F<sup>hi</sup> vs Siglec-F<sup>low</sup> neutrophil DEG (Lung Adenocarcinoma model) | Single cell | 29191879 | -<br>(Pre-acquired DEG in Table) |
| Siglec-F<sup>hi</sup> vs Siglec-F<sup>low</sup> neutrophil DEG (DEP-induced asthma model) | Bulk | 34653517 | GSE158040 |
| Siglec-F<sup>hi</sup> vs Siglec-F<sup>low</sup> neutrophil DEG (Myocardial infarction model) | Single cell | 33525909 | GSE157244 |

**Table S2.**
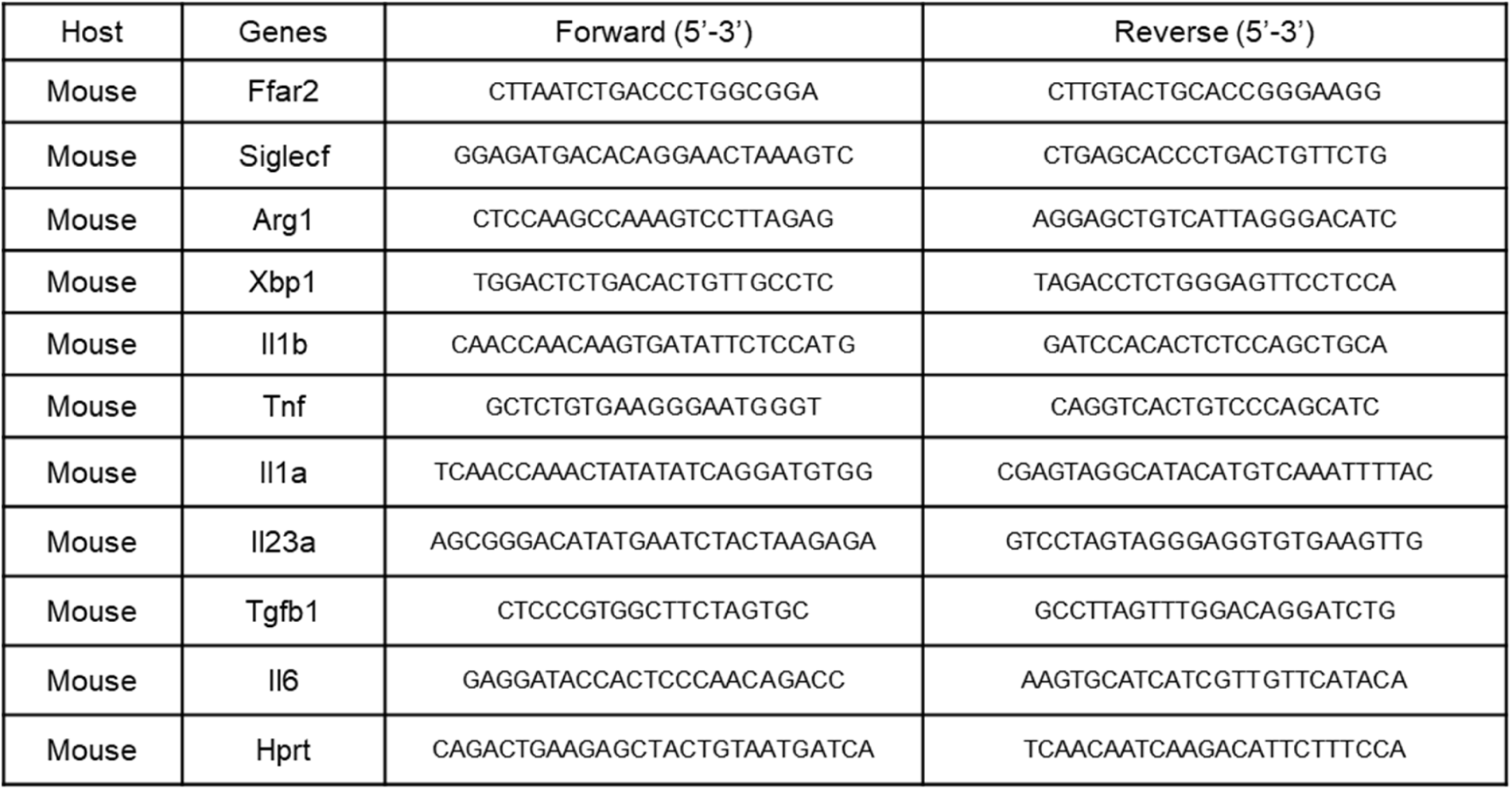
Primers for qPCR.

